# Turing pattern formation in reaction-cross-diffusion systems with a bilayer geometry

**DOI:** 10.1101/2023.05.30.542795

**Authors:** Antoine Diez, Andrew L. Krause, Philip K. Maini, Eamonn A. Gaffney, Sungrim Seirin-Lee

## Abstract

Conditions for self-organisation via Turing’s mechanism in biological systems represented by reaction-diffusion or reaction-cross-diffusion models have been extensively studied. Nonetheless, the impact of tissue stratification in such systems is under-explored, despite its ubiquity in the context of a thin epithelium overlying connective tissue, for instance the epidermis and underlying dermal mesenchyme of embryonic skin. In particular, each layer can be subject to extensively different biochemical reactions and transport processes, with chemotaxis – a special case of cross-diffusion – often present in the mesenchyme, contrasting the solely molecular transport typically found in the epidermal layer. We study Turing patterning conditions for a class of reaction-cross-diffusion systems in bilayered regions, with a thin upper layer and coupled by a linear transport law. In particular, the role of differential transport through the interface is explored together with the presence of asymmetry between the homogeneous equilibria of the two layers. A linear stability analysis is carried out around a spatially homogeneous equilibrium state in the asymptotic limit of weak and strong coupling strengths, where quantitative approximations of the bifurcation curve can be computed. Our theoretical findings, for an arbitrary number of reacting species, reveal quantitative Turing conditions, highlighting when the coupling mechanism between the layered regions can either trigger patterning or stabilize a homogeneous equilibrium regardless of the independent patterning state of each layer. We support our theoretical results through direct numerical simulations, and provide an open source code to explore such systems further.

## 1 Introduction

### 1.1 Biological motivation and study objectives

Among many other pioneering works, Turing [63] introduced a new, mathematical way to understand symmetry-breaking phenomena in biology. Over the intervening decades, his concept of a diffusion driven instability in reaction-diffusion systems has been extensively studied in the mathematical literature and confronted with biological observations and experiments [31, 4, 44, 45]. Despite extensive evidence that Turing patterning mechanisms can explain numerous complex phenomena observed in nature, a core challenge is modernizing Turing’s ideas to accommodate advances in our understanding of the living world that have emerged since his study. In its most classical version, Turing’s theory predicts that a homogeneous equilibrium state between two species can be broken by the sole effect of the increased diffusion of one species [45, Chapter 2], as in the renowned local activation/long-range inhibition paradigm advocated in [23, 40]. Many recent works have refined this theory and proposed more realistic biological scenarios, in particular regarding the geometry of the domain [32], the complexity of the signalling network [34] or the inclusion of other influences, such as cross-diffusion [18, 37], mechanical forces or active cell transport phenomena [46, 38, 64]. Contemporary perspectives on Turing systems are summarized in [32].

One refinement of Turing’s ideas that has been under-explored is the bilayer structure of many biological pattern forming systems. For instance, layered development has been indicated as relevant for the morphogenesis of the plant shoot apical meristem [20], cell-membrane Turing self-organisation modulated by signalling molecules in the cytosol [35], cell-cell membrane communication processes [61], and propagation problems in population ecology [21, 58, 7]. Outside biology, bilayer systems have been studied in chemistry in the context of the so-called CIMA and CDIMA experiments [5] which have motivated many analytical and experimental works on the patterning mechanisms of bilayer systems [65, 66, 67, 6, 9].

In particular, an important biological system that especially motivates this work is the embryonic skin, which consists of a thin epidermal layer superimposed on the extracellular matrix of the dermal mesenchyme. While the former is a thin layer mostly composed of tightly packed cells with limited movement, the latter can be much deeper and is composed of a network of collagen fibers supporting motile cells. During embryonic development, the interplay between these two layers gives rise to various repeating anatomical patterns such as hair follicles [25] (Fig. 1a), feather placodes [28] or finger prints [24]. These studies have identified complex signalling networks of diffusive molecules produced in each layer and which interact with the motile mesenchymal cells in order to initiate local cell clustering. Although mathematical reaction-diffusion-chemotaxis models [3, 49, 19] have been able to accurately reproduce the biological observations, they have focused on a simplified mono-layer geometry, i.e. a single domain in which all the species interact. However, biological experiments in [28] have stressed the importance of the bilayer structure on pattern formation by considering chimera skins composed of an epidermis from one species and a dermis from another. Depending on the species considered, the patterning ability may be conserved, or not, as well as the periodic structure of the patterning. This demonstrates that the coupling between the two layers is in itself a crucial component for symmetry-breaking.

**Figure 1:**
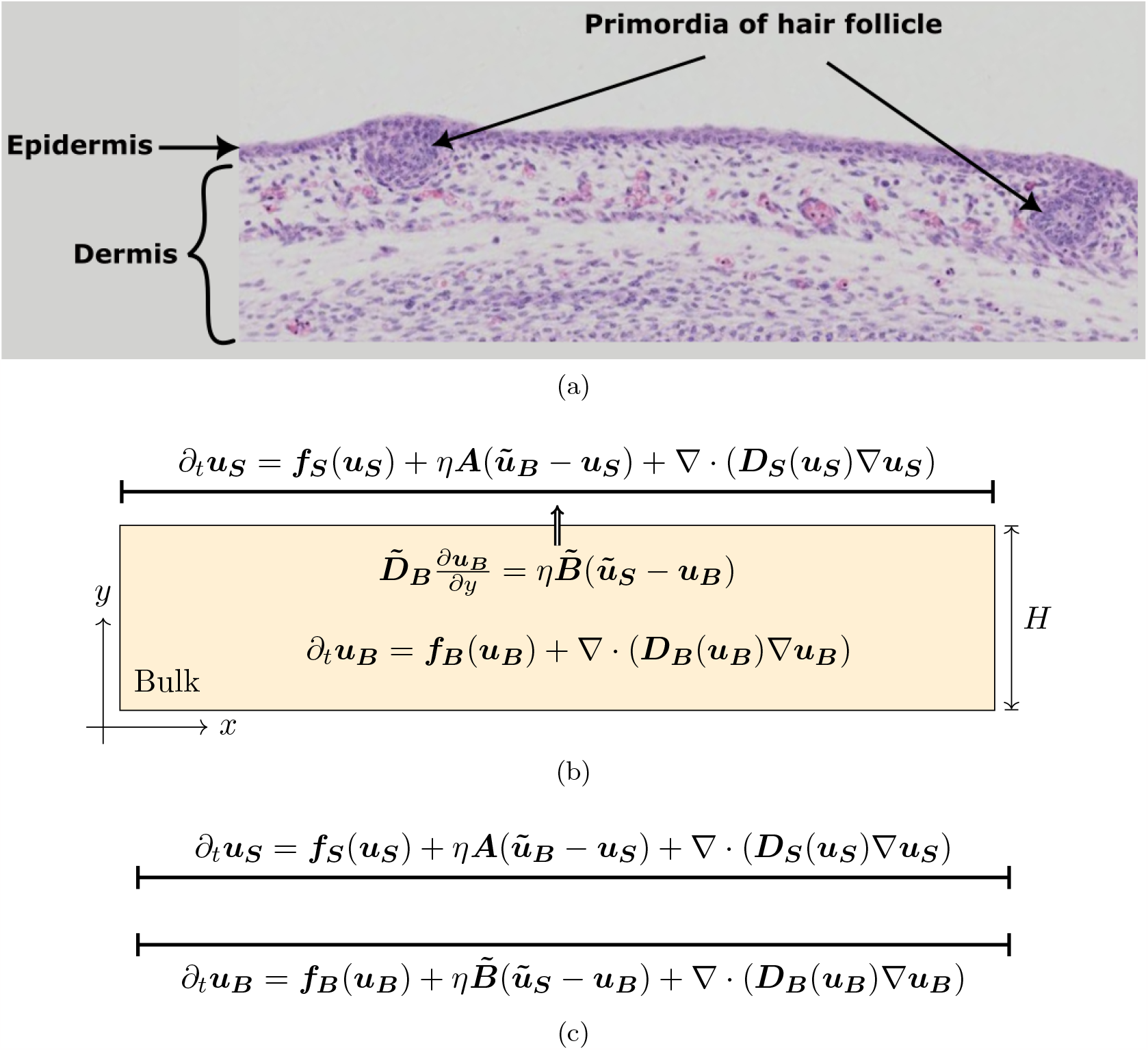
Embryonic skin patterning in bilayer tissue and domain geometry. (a) An example of skin patterning: the emergence of primordia of hair follicles at the interface between the epidermis and the dermis of a mouse embryo at 15.5 days post coitum. The formation of these condensates of mesenchymal cells can be explained by a complex network of reaction-diffusion and chemotaxis interactions between the two layers (see [25, 28]). Original picture from the e-Mouse Atlas Project under a CC BY 3.0 licence [2]. (b) Mathematical model in a 1D-2D geometry. The epidermis (later referred to as the surface) is modelled by a one-dimensional layer while the dermis (later referred to as the bulk) is two-dimensional to take into account its depth. The arrow symbolizes the boundary condition along the outward normal of the domain. (c) Mathematical model in the 1D-1D geometry where the two layers are taken to be one-dimensional.

Hence, on noting that cross-diffusion encompasses chemotaxis as a special case, the focus of this study will be the mathematical derivation of patterning conditions for bilayered reaction-cross diffusion systems, coupled via linear transport between the layers. In particular, our objective will be to determine when the bilayer structure, with a thin upper layer represented via a one dimensional domain, is predicted to enhance self-organisation or stabilise the homogeneous equilibrium of the system.

### 1.2 Related works and previous theoretical results

Bilayer systems similar to, but simpler than, the ones considered in the present article have been analytically and computationally studied, in particular in [33] and in [9]. The former article [33] is motivated by patterns formed by bacteria growing on an agar substrate and thus considers a pure reaction-diffusion system with a passive bulk (i.e. with only diffusion). Depending on the thickness of the layers, several asymptotic instability conditions and reduced dispersion relations are derived and studied numerically. The latter article [9] provides a detailed analysis of a coupled reaction-diffusion system of two exactly identical two-component layers. In this setting, the linear stability analysis of the coupled system is amenable to block-matrix computations which reduce the problem to a classical 2D eigenvalue problem and thus allows for a detailed bifurcation analysis.

Much earlier works were also motivated by the role of reaction-diffusion mechanisms in skin patterning problems. In particular, in [48], a very similar example based on a two-component reaction-diffusion system is studied numerically with the aim of distinguishing the mechanisms leading to spot and stripe patterns. The authors assume equal reaction terms in both layers but possibly different diffusion coefficients and a nonlinear transport law between the two layers. Patterning is assumed to be driven by the epidermis where the homogeneous state is unstable. Related works [60, 11, 46] have also considered mechano-chemical models of pattern formation in the skin, with an overview presented in [45, Chapter 6].

Other closely related theoretical works [56, 55, 35, 36, 26, 50, 51] in the literature are motivated by the cell membrane-cytosol system and thus focus on a spherical geometry and most often a passive bulk. Although in this context the curvature of the domain itself may have an important effect on pattern formation, we restrict ourselves to a planar geometry which is more relevant for the example of skin patterning. Another class of related models recently studied in the literature are the so-called compartmental models [53, 52] where two or more reaction-diffusion compartments are spatially coupled through a passive diffusive bulk or an interface [62]. In our setting, we do not consider a passive bulk, since chemotaxis and other nonlinear reaction-diffusion effects are anticipated also in the bulk (for instance again in the case of skin patterning). We will however only consider homogeneous equilibria in each layer, which is in contrast with the recent articles [50, 51] where the passive (linear) bulk in a spherical geometry allows for the derivation and systematic study of heterogeneous bulk equilibria. Note also that the existence of, possibly non-equal, homogeneous equilibria in each layer with arbitrary reaction kinetics cannot always be guaranteed so we will discuss and introduce an appropriate modelling framework where this case can be considered.

Although all these works study patterning conditions for various types of coupling between reaction-diffusion systems, their modelling frameworks are quite different from the bilayer structure that we consider here. In this article, we will consider a more general cross-diffusion framework (which includes chemotaxis models) with two non-necessarily identical active layers and an arbitrary number of interacting species. We give a set of quantitative conditions for (non)-patterning in the asymptotic limit of small and large couplings and provide several examples of patterning scenarios, theoretically, and with numerical evidence for reaction-diffusion and chemotaxis systems. In particular we consider the general scenario of pattern formation driven by an individual layer or their mutual coupling. Our analysis is based on classical linear stability analysis in the context of multi-component reaction-cross-diffusion systems. Since explicit analytical results cannot typically be obtained in this situation, we derive patterning conditions via quantitative approximations of the bifurcation curve depending on the coupling strength. In the weak coupling case, related perturbation techniques have been used in a different context for the study of weakly coupled oscillators and reaction-diffusion networks [15, 16].

The present article is structured as follows. The modelling framework is described in Section 2 and the main contributions are summarized in Section 2.4. As a starting point of the analysis, the dispersion relations and bifurcation conditions are written in full generality in Section 3. We then split the analysis into two asymptotic cases, first the weak coupling case in Section 4 and secondly the large coupling case in Section 5. Phenomena in the intermediate coupling case are briefly described in Section 6 before the conclusions and discussion in Section 7. The supplementary material contains a description of the numerical methods (Appendix A) and the list and description of the supplementary videos (Appendix B).

## 2 Models and basic properties

### 2.1 1D surface - 2D bulk space model

We first consider a suitably non-dimensionalised bilayer system Ω_*S*_ *∪* Ω_*B*_ where the thin upper layer of length *L >* 0, Ω_*S*_ = [0, *L*] – referred to as the surface – is taken to be infinitesimally thin compared to the lower layer Ω_*B*_ = [0, *L*] *×* [0, *H*] – referred to as the bulk – which has a depth *H >* 0 (Fig. 1b). We consider *n* species in the surface and *m* species in the bulk. In both layers we assume interactions of these species according to a system of reaction-cross-diffusion equations. Their concentrations are denoted by^1^ ***u***_***S***_(*x*) *∈* ℝ^*n*^ in the surface at the location *x ∈* Ω_*S*_ and ***u***_***B***_(*x, y*) *∈* ℝ^*m*^ in the bulk at the location (*x, y*) *∈* Ω_*B*_. We will always consider *m ≥ n* and we assume that the *n* species in the surface can diffuse to the bulk and vice-versa for the first *n* components of the bulk species ***u***_***B***_. There are *m− n* species which do not diffuse to the surface and remain in the bulk. The concentrations of these *m − n* species are given by the last *m − n* components of the vector ***u***_***B***_ *∈* ℝ^*m*^. The mathematical analysis would hold similarly in the reverse case but we choose *m ≥ n* for both parsimony and also noting the biological case of skin patterning where an extra chemotaxis component would be included in the bulk domain. Then, the general form of the model is given by

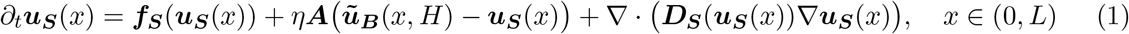

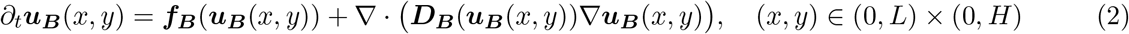

where the matrix ***A*** *∈* ℝ^*n×n*^ specifies the exchange rates of the different species between the two layers and ***ũ***_***B***_ *∈* ℝ^*n*^ denotes the vector constructed by taking the first *n* components of ***u***_***B***_ *∈* ℝ^*m*^, that is, when ***u***_***B***_ = (*u*_*B*,1_, …, *u*_*B,m*_)^T^ *∈* ℝ^*m*^,

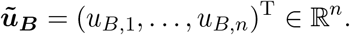

where the superscript T denotes the transpose. The parameter *η ≥* 0 represents the strength of the coupling between the two layers. The reaction functions ***f***_***S***_ and ***f***_***B***_ are arbitrary and the crossdiffusion matrices ***D***_***S***_ and ***D***_***B***_ are positive definite.

On the lateral sides and the bottom side of the bulk, we assume zero-flux boundary conditions. At *y* = *H*, we consider the following linear transport law for the bulk species:

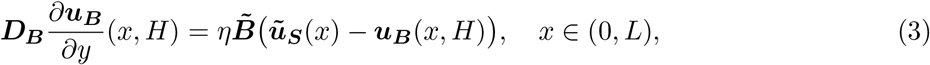

with 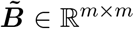 denoting the matrix constructed on padding ***B*** *∈* ℝ^*n×n*^ by zeros, where ***B*** specifies linear transport from the bulk to the surface. Similarly, ***ũ***_***S***_ *∈* ℝ^*m*^ denotes the vector constructed from ***u***_***S***_ *∈* ℝ^*n*^ by adding *m − n* rows with zero components, that is

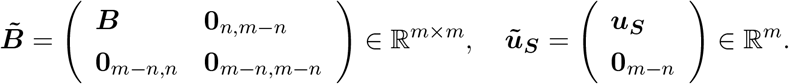

Furthermore, note that ***B*** *−* ***A*** corresponds to interfacial sources/sinks of species, which are allowed for generality, but these will often be zero in applications.

Note that as in earlier works [48], one could also consider nonlinear transport laws but for simplicity and because we will only carry out a linear stability analysis that would also linearize this part of the equation, we will only work with linear transport between the layers.

As motivated in the introduction, we focus on a model representing the bilayer structure of embryonic skin [25, 28, 24], with bulk chemotaxis where cells restricted to the bulk are chemo-attracted by *n* species, which diffuse between the two layers (Fig. 1). In this case *m* = *n* + 1 and ***u***_***B***_ = (***ũ***_***B***_, *c*) where *c*(*t, x, y*) denotes the cell concentration. Assuming a classical reaction-diffusion interaction for the chemical species, the equation in the bulk reduces to

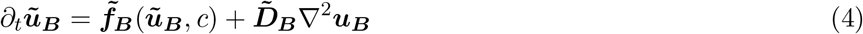

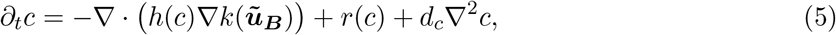

where 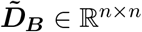 is a positive definite diagonal matrix and *d*_*c*_ *>* 0. The representative examples for *h*(*c*), *r*(*c*) and *k*(***ũ***) are given by

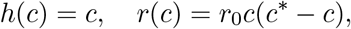

with *r*_0_, *c*^*\**^ positive constants, and *k*(***ũ***) = *u*_1_, where *u*_1_ is the first component of ***ũ*** *∈* ℝ^*n*^. The boundary condition at the interface can be rewritten

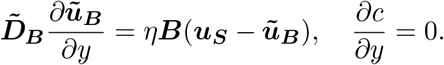

This corresponds to Eqs. (2), (3) with

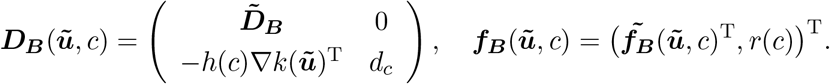

### 2.2 1D surface - 1D thin bulk space model

In the embryonic skin example described in the introduction, local cell clustering is typically observed only at the interface between the epidermal and dermal layers (Fig. 1a) and the whole process takes place on a spatial range which does not exceed the diameter of a few cells [25, 28]. Thus, we will also consider the case of an infinitesimally thin bulk (Fig. 1c).

Under an appropriate rescaling of the coupling intensity *η*, the 1D-2D model (1)-(3) can be reduced to two coupled one-dimensional equations. In particular, let us consider a bulk concentration ***u***_***B***_ and a bulk depth of *H ≡ ε >* 0, with *ε ≪* 1 corresponding to the additional assumption that the bulk depth is much smaller than any other lengthscale in the 1D-2D model of the previous subsection. We then introduce the rescaled concentration 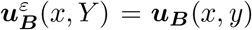 where *Y* = *ε*^*−1*^*y ∈* [0, 1]. With this change of variable, Eq. (2) becomes

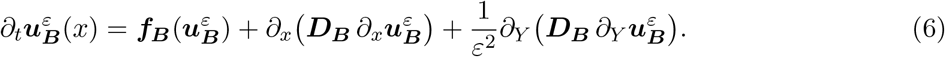

We also rewrite 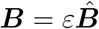, with the components of 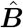 scaling as *O*(1), to ensure there is a balance between the inter-layer flux and reaction terms of the rescaled bulk equations, noting this scaling will also be inherited by 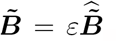. In particular, for other scalings, after the limit of infinitesimal bulk thickness has been taken, only one of the reaction term or the inter-layer flux will appear in the leading order dominant balance. If the reaction terms dominate, then the layers are uncoupled and the patterning is based on the individual layer dynamics at leading order, while if the inter-layer transport dominates then there is no chemotaxis or interaction between the signalling molecules, which again is not the system that is to be considered, as biological mechanisms of self-organisation are lost. Thus, with the scaling 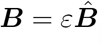, the boundary condition Eq. (3) becomes:

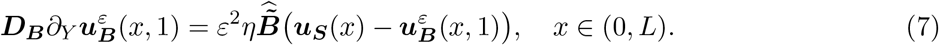

We then expand 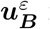 in powers of *ε*,

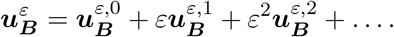

Carrying out this expansion in Eq. (6), we obtain at order *ε*^*−1*^ and *ε*^*−*2^

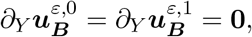

and at order 0, we have

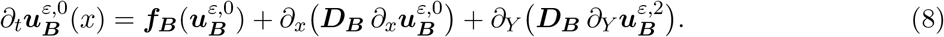

For the expansion in the boundary condition Eq. (7), we obtain

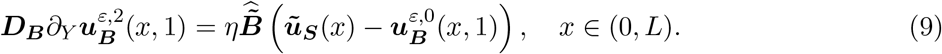

Thus, noting that 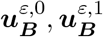 are independent of *Y*, integrating equation Eq. (8) in *Y* between 0 and 1, using the boundary condition (9) and, for notational simplicity, dropping the hat of 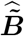 and the super-script label “*ε*, 0”, results in

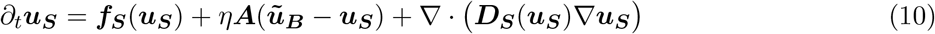

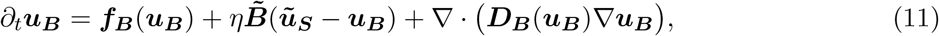

where all the functions are evaluated at a point *x ∈* (0, *L*) and the spatial derivative is one-dimensional and taken along the *x* direction. Hence one may also observe that once the infinitesimal bulk thickness limit has been taken, and with the constraint ***A*** = ***B***, (where we recall that ***B*** is the *n× n* block of 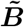 corresponding to species that can transport between the upper and lower regions), then there are no additional sources or sinks at the layer interface for the leading order equations. So the 1D-1D model is in some sense a special case of the 1D-2D model given in Eqs. (1)-(3), though with the important difference that the two layers of the 1D-1D model may not admit the same spatially-homogeneous equilibria, as explained in the following section.

### 2.3 Equilibria structure

Throughout this article, we assume that there exists a constant equilibrium point 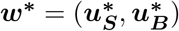 such that in the 1D-1D case

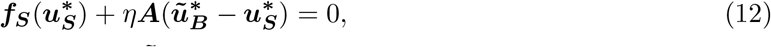

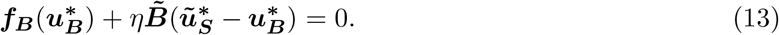

Note that this equilibrium point ***w***^***\****^ *≡* ***w***^***\****^(*η*) may depend on *η* and that 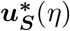 and 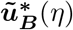 may be different from each other. As we are going to show in the following, the ability of the 1D-1D system to exhibit asymmetric equilibrium concentrations in the surface and the bulk can have a strong influence on pattern formation.

In the 1D-2D case, the boundary condition at the interface imposes 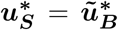 and ***w***^***\****^ is thus independent of the coupling strength *η* and only depends on the reaction terms, which must satisfy

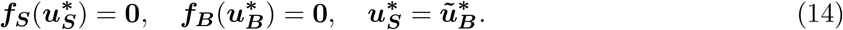

This condition can naturally be satisfied when ***f***_***S***_ and ***f***_***B***_ are proportional to each other (including when one reaction function is identically equal to zero). However, Eq. (14) does not impose any particular form for the reaction functions, which in principle could be completely different. In practice it may however require some ad hoc fine tuning of the parameters to ensure that the two reaction functions share the same equilibrium point. Note also that when *m > n*, the *m − n* pure bulk species concentrations may act as free parameters in the second relation. For instance in the chemotaxis case, the equilibrium cell concentration *c*^*\**^ can act as a parameter of the system, specified by a growth term of the form *r*(*c*) = *c*(*c*^*\**^ *− c*).

An important point to note is that the existence or uniqueness of such an equilibrium point in the 1D-1D case cannot be guaranteed by the sole intrinsic properties of the reaction terms, due to the dependence on *η*. For a sufficiently small coupling strength *η*, we will thus adopt a perturbative approach. We always assume that at *η* = 0, the uncoupled systems have a homogeneous equilibrium point. Then, by the implicit function theorem, for *η* sufficiently small, it is possible to find a curve *η* ↦ ***w***^***\****^(*η*) which satisfies the relations Eqs. (12)-(13), provided that the Jacobian matrices of ***f***_***S***_ and ***f***_***B***_ evaluated, respectively, at 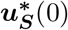 and 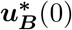 are invertible (see Section 4.3 for more details).

We also mention that we do not consider spatially heterogeneous equilibria in the 2D bulk as was done for instance in [50, 51] in a spherical geometry. In these articles, the derivation of such equilibria exploits the solvability of linear reaction-diffusion problems but we cannot expect to extend this derivation to arbitrary nonlinear bulk reaction kinetics. We thus focus on a perturbation analysis around a global homogeneous equilibrium (when it exists). However, we would like to stress that the 1D-1D model given by Eqs. (10)-(11) can be considered and analyzed independently since it has a very specific structure which potentially allows two asymmetric homogeneous equilibria, which is never possible for 1D-2D models.

### 2.4 Objectives and contributions

The goal of this study is to understand how Turing self-organisation is affected by the coupling in a bilayer geometry and to derive mathematical conditions for the linear instability of a homogeneous equilibrium in order to generate patterns. Thus we consider the fundamental mechanism of Turing pattern formation theory, namely diffusion-driven instability. We will focus our analysis on the two asymptotic regimes *η ≪* 1 and *η ≫* 1 and we will obtain explicit quantitative patterning conditions that can be numerically computed. We will always assume that the two layers independently can generate Turing patterns in general, though not necessarily with the parameters investigated, so that our main objective will be to determine analytically the cases where the coupling enhances or diminishes this patterning ability. Before proceeding to the mathematical analysis, we briefly summarize our main contributions below:

- **Weak coupling case (***η ≪* 1**) for the 1D-1D system:** Turing patterns can be formed by coupling two independently non-patterning layers provided that the exchange rate of one species is large enough (Section 4.2, Fig. 2b, Video 1) or that the equilibria of the two uncoupled layers are different (Section 4.3., Fig. 3).

**Figure 2:**
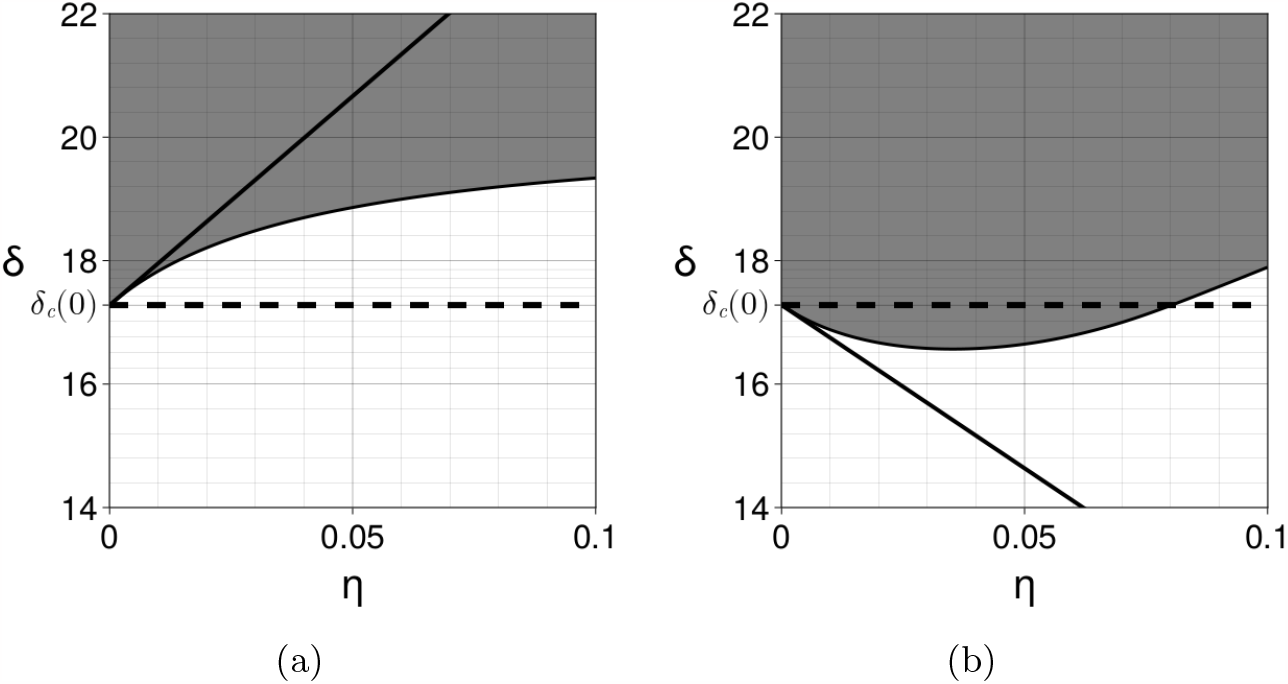
Coupling two identical Schnakenberg systems with *a* = 02305, *b* = 0.7695 and *s* = 1. The bifurcation parameter *δ* is the diffusion coefficient of the *v*-species. The instability region in the (*η, δ*) plane is depicted in grey and is computed numerically by applying the Routh-Hurwitz criterion to the fourth order polynomial Eq. (19) for the modes *k*_*q*_ = *qπ/L* with *L* = 1000 and *q ∈ {*0, …, 1500*}*. The thin solid line at the boundary of the instability region is computed by solving numerically Eqs. (20)-(21). The dashed horizontal line indicates the critical bifurcation parameter *δ*_*c*_(0) at *η* = 0. The thick solid line is the first-order approximation 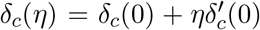 computed using Eq. (33). (a) When *α* = *β* = 1, the coupling degrades the ability to form Turing patterns in the sense that a higher diffusion coefficient is needed (i.e. the instability region is reduced when *η* is increased). (b) When *α* = 1 and *β* = 30, the coupling enlarges the parameter space in which Turing patterns are formed. See also Video 1. In both cases the bulk layer is in a non-patterning state with diffusion coefficients 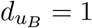 and 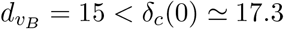.

**Figure 3:**
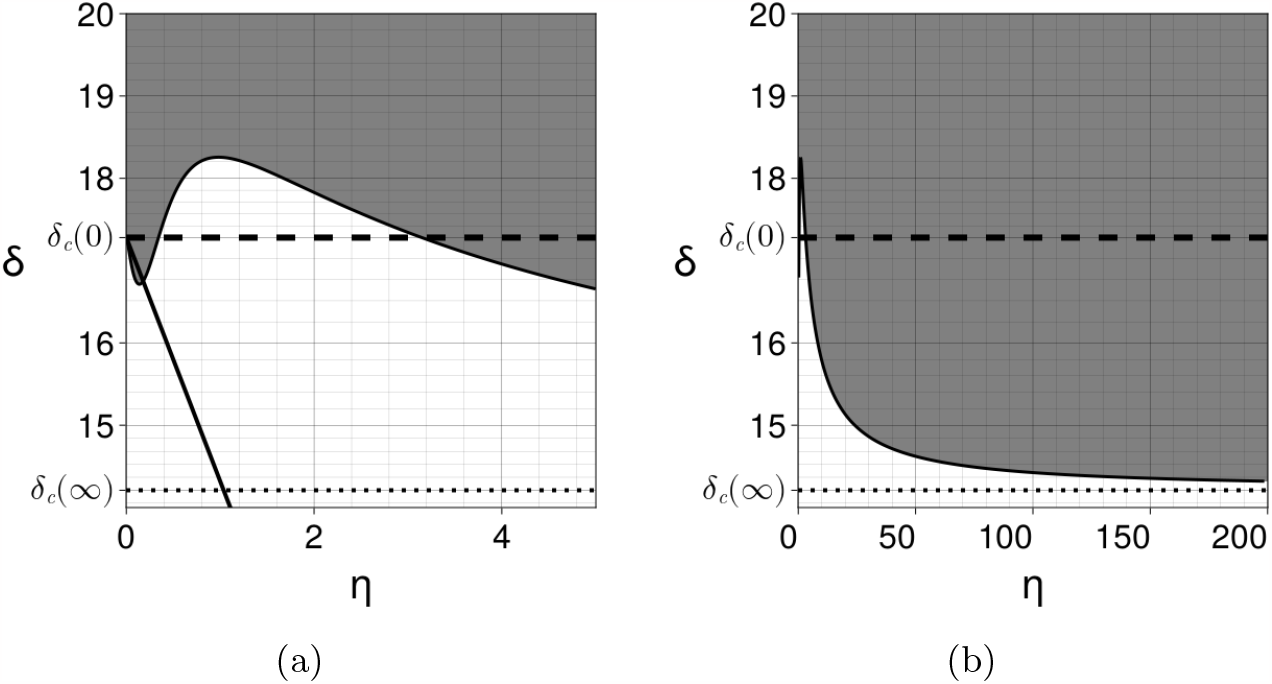
Coupling two different Schnakenberg systems with parameters ***A*** = ***B*** = ***I***, *a* = 0.15, *b* = 0.2, *s* = 0.5, 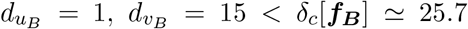, for the bulk (non-patterning state) and *a* = 0.2305, *b* = 0.7695, *s* = 2 for the surface. The instability region in grey and its boundary are computed numerically as in Fig. 2. The coupling enhances patterning for small and large *η* but reduces patterning for intermediate values. The first-order approximation at *η* = 0 is computed using Eq. (28). The asymptotic value is indicated by the dotted line and is computed using Eq. (42). (a) *η ∈* (0, 5). (b) *η ∈* (0, 200).
- **Weak coupling case (***η ≪* 1**) for the 1D-2D system:** we extend the 1D-1D results by studying (asymptotically) the influence of the bulk depth (Section 4.4.1). We also show different patterning scenarios in Section 4.4.3. (Videos 2-3-4).
- **Strong coupling case (***η ≫* 1**) for the 1D-1D system:** we show that the coupled system reduces to one single cross-diffusion equation (Section 5.1.) and find a simple criterion to determine whether the coupling stabilizes the homogeneous state (Section 5.2.1, Video 5) or enhances patterning (Section 5.2.2, Video 6).
- **Intermediate coupling case:** In Section 6. we briefly comment on the wide variety of patterning scenarios, and the difficulty in studying them in any generality, beyond the asymptotic regimes above.

## 3 Dispersion relations and bifurcation points

As a starting point, we derive the dispersion and bifurcation relations associated with a weak linear perturbation of the homogeneous state. In addition, in the 1D-2D case, the change of variable *y*^*′*^ = *H − y* is convenient, so as to locate the interface at *y*^*′*^ = 0, though the prime is dropped below.

### 3.1 1D-1D

For the 1D-1D system (10)-(11), taking into account the Neumann boundary conditions on the lateral sides, we consider a linear perturbation of the equilibrium point of the form

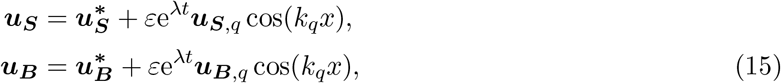

where *q* is an integer and *k*_*q*_ = *qπ/L*. Linearizing Eqs. (10)-(11) around this equilibrium point shows that at order *ε*, the perturbation vector 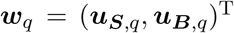 belongs to the kernel of the following matrix

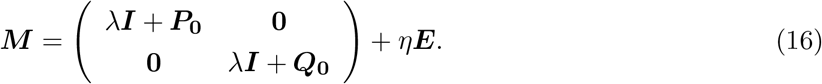

Here ***I*** is the identity matrix, 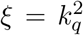 is treated as a continuous variable, ***P***_**0**_ = *ξ****D***_***S***_ *−* ***J***_***S***_, ***Q***_**0**_ = *ξ****D***_***B***_ *−* ***J***_***B***_, where ***J***_***B***_ and ***J***_***S***_ are the Jacobian matrices of ***f***_***S***_ and ***f***_***B***_ with respect to ***u***_***S***_ and ***u***_***B***_, respectively and,

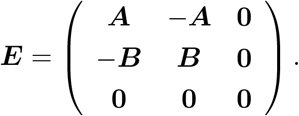

Unless specified otherwise, all expressions are evaluated at the equilibrium point ***w***^***\****^. Note again that ***w***^***\****^ may depend on *η* if the surface and bulk equilibrium concentrations are not equal.

The dispersion relation is given by the determinant of ***M*** being zero, i.e.

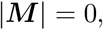

which is a polynomial relation in *λ* and *ξ* but possibly non polynomial in *η* due to the unknown dependence of ***w***^***\****^ on *η*. For a given set of parameters and a given *η*, it is nevertheless possible to determine the stability of each mode *ξ* by applying the Routh-Hurwitz criterion to the polynomial in *λ* of degree (*m* + *n*).

### 3.2 1D-2D

In the 1D-2D case, the amplitude of the perturbation in the bulk depends on the *y*-variable so that the linearized equations become

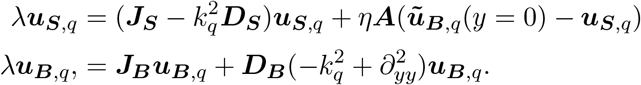

The second equation can be rewritten

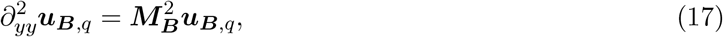

where the matrix ***M***_***B***_ is a square root of

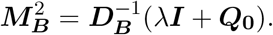

Note that in principle the square root of a matrix is not unique and may not even exist. The square root of a diagonal matrix can be constructed by taking the principal square root of the diagonal elements. This construction readily extends to diagonalizable matrices which form a dense open set. However, as we shall see, this particular convention does not play any role in the following. Solving Eq. (17) in *y*, with *y* = 0 corresponding to the interface, and no-flux conditions at *y* = *H*, gives

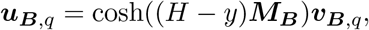

for some vector 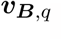, with the interfacial conditions still to be imposed. These, in turn, give

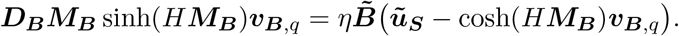

In the previous expressions, the hyperbolic sine and cosine of a matrix are defined by their power series expansion, namely for a matrix ***M***,

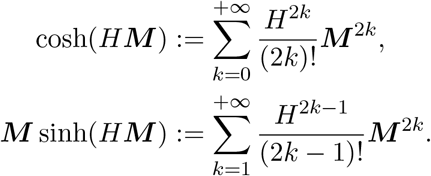

Note in particular that these are functions of ***M*** ^2^, so that the final results do not depend on taking a matrix square root, nor the choice of the square root.

Finally, we conclude that 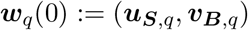 belongs to the kernel of the following matrix

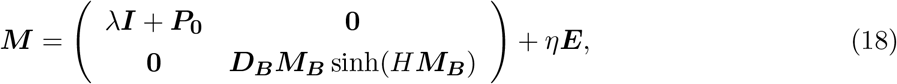

with

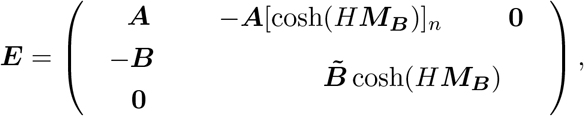

where [cosh(*H****M***_***B***_)]_*n*_ denotes the first *n* rows of cosh(*H****M***_***B***_). Note that, unlike *λ****I*** + ***P***_**0**_, the matrix ***M***_***B***_ does not depend linearly on *λ* or *ξ* due to the square root and the hyperbolic sine and cosine. Consequently, the dispersion relation

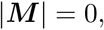

is a transcendental relation in all variables. In particular, there is no standard criterion, like the Routh-Hurwitz criterion in the polynomial case, to determine the stability of a given mode. Note also that this dispersion relation is a straightforward extension of the model studied in [33] in a 2D-2D reaction-diffusion scenario with a passive bulk layer.

As an example, in the chemotaxis case given by Eqs. (4)-(5), one can check that

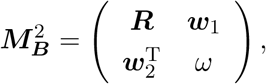

with

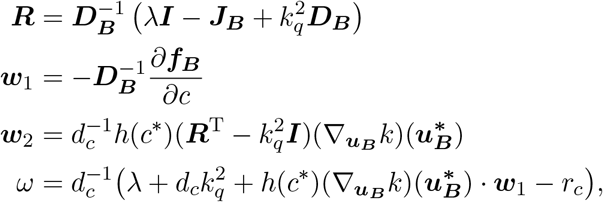

where *r*_*c*_ = *∂r/∂c*, evaluated at the equilibrium point.

### 3.3 Bifurcation points

In the following, all the parameters are assumed to be fixed except for one, generically denoted by *δ* which will be taken as a bifurcation parameter. Classically, *δ* is the diffusion coefficient of one of the species, for instance the inhibitor diffusion in a classical two-species reaction-diffusion system [45, Chapter 2]. Note however that in the following, the parameter *δ* could be any parameter of the model, such as the chemotaxis strength. Following the classical Turing theory for marginal instability, for a given *η* we are interested in the critical value *δ*_*c*_ of the bifurcation parameter at which *λ* = 0 is a solution of the dispersion relation, while all the other solutions have a negative real part (i.e. a Turing bifurcation). In this article, we will only consider the case of a Turing bifurcation and leave the case of Hopf (or Wave) bifurcations (i.e. associated with *λ* = *iρ, ρ* real) for future work. The goal of this section is to find a set of algebraic relations on (*ξ, δ*) that should be satisfied for a Turing bifurcation to occur.

Highlighting the dependence with respect to the various parameters, the dispersion relation reads

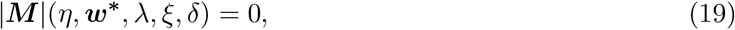

where ***M*** is given by Eq. (16) or Eq. (18). Thus, for a given *η* and a given equilibrium ***w***^***\****^, defining the function (*ξ, δ*) *↦ a*_0_(*η*, ***w***^***\****^, *ξ, δ*) ≔ |***M*** |(*η*, ***w***^***\****^, 0, *ξ, δ*), the critical values *ξ*_*c*_ and *δ*_*c*_ at which a Turing bifurcation occur satisfy the following relation:

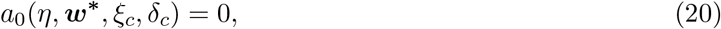

and our goal is to study the dependence of this solution on *η* and ***w***^***\****^. More precisely, when *η* = 0 (i.e. when the two layers are independent), we will assume that a Turing bifurcation occurs for one of the layers when the bifurcation parameter crosses a value *δ*_*c*_(0). Then, when *η >* 0, one of our main goals will be to compare the critical value *δ*_*c*_(*η*) of the bifurcation parameter of the coupled system with the critical value *δ*_*c*_(0) of the uncoupled system. More generally, we will give quantitative estimates on how *δ*_*c*_(*η*) behaves as *η* increases.

Note also that since all the other solutions of the dispersion relation are assumed to have a non positive real part, it follows that *a*_0_ *≥* 0 in a *ξ* neighbourhood of *ξ*_*c*_, which imposes the second relation

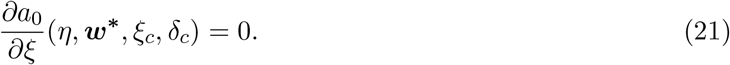

In the following, it will be useful to expand *a*_0_ as

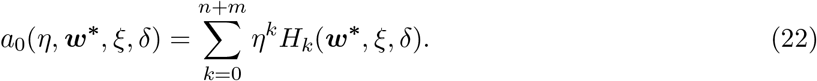

The first term is computed by setting *η* = 0, in which case Eq. (20) reduces to a block-diagonal determinant:

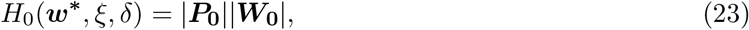

where ***P***_**0**_ = *ξ****D***_***S***_ *−* ***J***_***S***_ and ***W***_**0**_ = ***Q***_**0**_ = *ξ****D***_***B***_ *−* ***J***_***B***_ in the 1D-1D case. For the 1D-2D case, ***P***_**0**_ has the same definition but 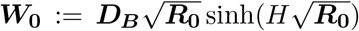 with 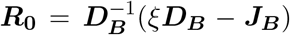 and 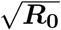 denoting any square root of ***R***_**0**_. For the second term, we recall Jacobi’s formula for the differential of the determinant function,

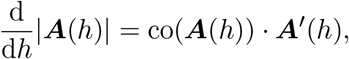

for any matrix-valued differentiable curve *h ∈* ℝ ↦ ***A***(*h*), with derivative at *h* denoted by ***A***^*′*^(*h*), where ***A*** *·* ***B*** ≔ Tr(***A***^T^***B***) denotes the usual matrix dot product and co(***A***) is the comatrix of the matrix ***A*** (which is the transpose of the adjugate matrix).

Consequently, we obtain that

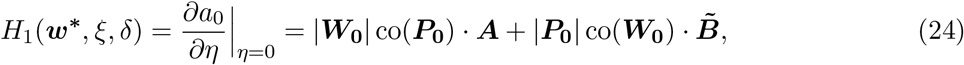

in the 1D-1D case and

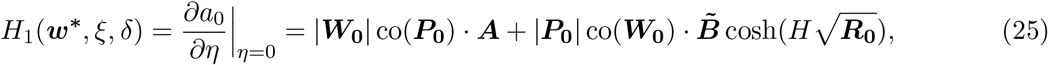

in the 1D-2D case.

**Remark 1**. *The expansion* (22) *will be useful to study the small coupling case η ≪* 1 *in the next section. Following the same methodology, it would possible to compute the full Taylor expansion of a*_0_ *using the Faa di Bruno formula, although, due the algebraic complexity, it does not seem possible to obtain exploitable analytical results beyond the first order*.

## 4 Weak coupling case for the 1D-1D and 1D-2D models

When *η* = 0, the surface and bulk systems are decoupled and we assume that at least one of them has a bifurcation parameter with the ability to produce Turing patterns. Classically, it is possible to compute, explicitly or numerically, the critical value of this bifurcation parameter, *δ*_*c*_(0), as well as the critical wave number *ξ*_*c*_(0) (see for instance [45, Section 2.3.] as well as Section 4.2.2. in the case of a two-species reaction-diffusion system). The goal of this section is to study how this critical parameter changes when the two systems are coupled with a small coupling strength *η >* 0.

### 4.1 General formula

We first give a simple criterion for the existence of a critical bifurcation parameter function *η ↦ δ*_*c*_(*η*) and *η ↦ ξ*_*c*_(*η*) in a neighbourhood of *η* = 0. Since these critical values are defined by Eqs. (20)-(21), by the implicit function theorem and the expansion given in Eq. (22), this reduces to proving that

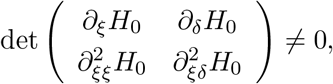

where all the partial derivatives are evaluated at (*ξ*_*c*_(0), *δ*_*c*_(0)). We will call surface-driven (resp. bulk-driven) symmetry-breaking the case where *δ* is the diffusion coefficient of a surface (resp. bulk) species and thus at *δ*_*c*_(0), |***P***_**0**_| = *∂*_*ξ*_|***P***_**0**_| = 0 and |***Q***_**0**_| 0 (resp. with ***Q***_**0**_ and ***P***_**0**_ switched). Without loss of generality, let us therefore assume a surface-driven symmetry breaking (again, otherwise ***P***_**0**_ and ***Q***_**0**_ are simply switched). In this case, using Eq. (23), the above condition thus reduces to

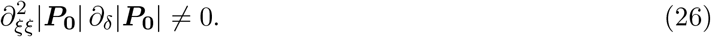

This condition is fulfilled, for instance, for a classical two-species activator-inhibitor system away from higher codimension points, as it is just a transversality condition. We will always assume that Eq. (26) holds in the following.

In order to know if the coupling increases or decreases the ability to form patterns, we want to compute the derivative of *δ*_*c*_ at *η* = 0 and thus obtain a first order approximation of the bifurcation curve *η* ↦ *δ*_*c*_(*η*). To do so, since

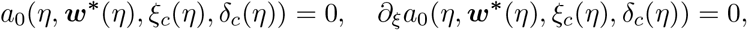

taking the derivative of these relations with respect to *η* (and using ^*′*^ to indicate derivatives with respect to *η*), we find that 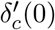 and 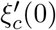 satisfy the following system of coupled partial differential equations

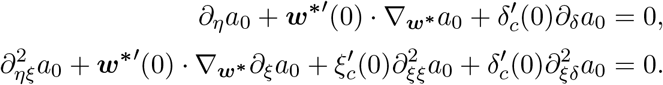

Using the expansion in Eq. (22), this reduces to

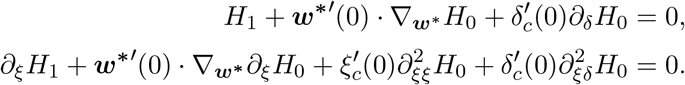

In particular, we deduce the general formula

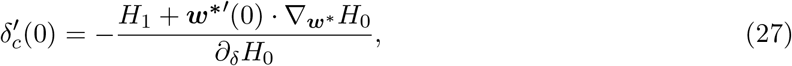

where *H*_1_ and *H*_0_ are evaluated at (***w***^***\****^(0), *ξ*_*c*_(0), *δ*_*c*_(0)). This formula can be further simplified by applying Eqs. (23)-(24) in the different cases that we will consider below. For later convenience, we summarize the results in the following straightforward proposition.

#### Proposition 1.

*Under the assumption that Eq*. (26) *holds true, the derivative at η* = 0 *of the bifurcation curve η* ↦ *δ*_*c*_(*η*) *is given by the following formulas*.

- ***In the 1D-1D surface-driven symmetry breaking case***,

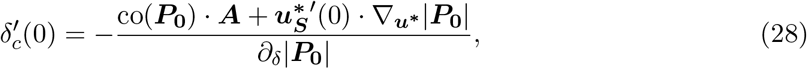
- ***In the 1D-1D bulk-driven symmetry breaking case***, *the same formula Eq*. (28) *holds with* 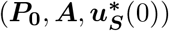 *replaced by* 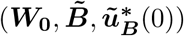.
- ***In the 1D-2D surface-driven symmetry breaking case***, *we recall that the two layers must share the same equilibria, so* ***w***^***\*****′*^(0) = 0 *and consequently*

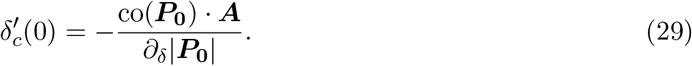
- ***In the 1D-2D bulk-driven symmetry breaking case***, *we need to take into account the additional hyperbolic cosine term in Eq*. (25), *and finally obtain*

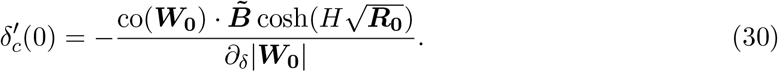

One surprising consequence of the formula Eq. (28) (and similarly for its variants Eqs. (29)-(30)) is that at first order when *η ≪* 1, the function *η ↦ δ*_*c*_(*η*) does not depend on the bulk (resp. surface) parameters. When 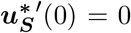, it is an intrinsic property of the surface (resp. bulk) system. When 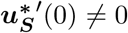, as explained in Section 4.3., there is an additional contribution, but this only depends on the difference between the equilibrium concentrations of the bulk and the surface.

We recall that at (*ξ*_*c*_(0), *δ*_*c*_(0)), |***P***_**0**_| = *∂*_*ξ*_|***P***_**0**_| = 0 and |***P***_**0**_| *≥* 0 elsewhere for surface driven symmetry breaking. Moreover, patterning occurs in the surface system as soon as there exist parameters (*ξ, δ*) such that |***P***_**0**_|(*ξ, δ*) *<* 0. Consequently, when *∂*_*δ*_|***P***_**0**_| *<* 0, patterning occurs in the surface layer independently for *δ > δ*_*c*_(0) and when *δ < δ*_*c*_(0) if *∂*_*δ*_|***P***_**0**_| *>* 0. The second case is typically encountered when *δ* is the diffusion parameter of the cells in a chemotaxis system, whereas the first case corresponds, for instance, to a classical activator-inhibitor system with *δ* the diffusion parameter of the inhibitor. Following this observation, one can conclude that the coupling enhances patterning when 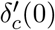 and *∂*_*δ*_|***P***_**0**_| have the same sign. For instance, when *∂*_*δ*_|***P***_**0**_| *<* 0, this would imply that *δ*_*c*_(*η*) *< δ*_*c*_(0) for *η* small enough. Due to Formula (28), this occurs if and only if

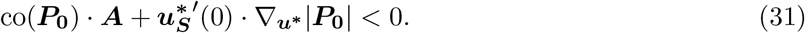

The increased or decreased ability to form Turing patterns thus depends on the balance between the two terms in Eq. (31). The second term may be non-zero only in the case where the two layers have different equilibrium concentrations. Note that this can only happen in the 1D-1D case. The contribution of this term is difficult to estimate in full generality, so we will give some examples in Section 4.3. . We now focus on the contribution of the first term, theoretically and for various examples.

### 4.2 Global equilibrium

In this section, we assume the following.

#### Assumption 1

(Global equilibrium). *The two layers share the same equilibrium when they are uncoupled, i*.*e*.

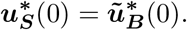

*In particular, this value remains a global homogeneous equilibrium for any η >* 0, *which implies that* 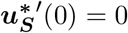.

#### 4.2.1 Theoretical considerations with equal surface and bulk equilibrium concentrations

Under Assumption 1, the second term in Eq. (31) vanishes and hence we focus on the contribution of the first term. That is, we want to compute the sign of

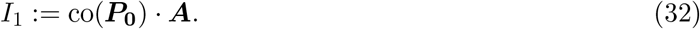

Note that in the 1D-1D model, the surface and bulk are exchangeable so the main result of this section, summarized in the following proposition, can be immediately translated to 1D-1D bulk-driven symmetry breaking. Moreover, owing to Eq. (29), the analysis in this section also applies to surfacedriven symmetry breaking in the 1D-2D case. The bulk-driven symmetry breaking situation in the 1D-2D model will be discussed in Section 4.4.1.

##### Proposition 2.

*Under Assumption 1 in the 1D-1D surface-driven symmetry breaking case, the following results hold*

- *if* ***A*** = ***I***, *then* 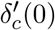 *and ∂*_*δ*_|***P***_0_| *have opposite signs (i*.*e. coupling degrades the ability to form patterns);*
- *if* ***D***_***S***_ *is diagonal with constant non-negative coefficients (pure reaction-diffusion system), then there exists a diagonal matrix* ***A*** *such that* 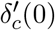 *and ∂*_*δ*_|***P***_0_| *have the same sign (i*.*e. coupling enhances patterning)*.

*Proof*. Let us first consider the case ***A*** = ***I***. The quantity (32) is thus equal to the sum of the *n* principal minors associated with the principal submatrices of order (*n −* 1). It can be shown that this quantity is equal to the product of the (*n −* 1) non-zero eigenvalues of ***P***_**0**_ [41, Section 7.1.]. By definition, at (*ξ*_*c*_(0), *δ*_*c*_(0)), all these eigenvalues have a positive real part and thus we conclude that *I*_1_ *≥* 0 and thus, using Eq. (28) and Assumption 1, *∂*_*δ*_|***P***_**0**_| and 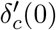 have opposite signs. This implies that the coupling always degrades the ability to form patterns. In order to enhance patterning, one needs to choose a matrix ***A*** whose elements corresponding to negative cofactors are non-negative and large. For pure reaction-diffusion systems – that is ***D***_***S***_ is diagonal with nonnegative coefficients – such cofactors always exist since

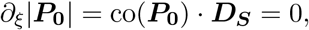

which implies that at least one diagonal cofactor is negative. □

It is important to note that since the first order approximation 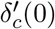 is directly proportional to the norm of ***A***, it can be made as small as possible by an appropriate scaling of the exchange rates. In theory, this fact thus allows the possibility that patterning can be enhanced by the layer-coupling given an arbitrarily small diffusion of the reactive species. Compared to the Turing theory for reaction-diffusion systems, the differential transport (i.e. the fact that one of the interacting species has a higher exchange rate through the interface) can compensate for, or even replace, the usually theoretically necessary differential diffusion for the classical version of the Turing mechanism, which has long been a subject of debate in real-world biological systems [4, 40, 37, 64].

In the following section, we will illustrate these phenomena through various examples in the 1D-1D situation.

#### 4.2.2 Two-species 1D-1D reaction-diffusion systems

For this example, we consider only two reacting species with concentrations

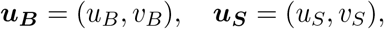

which share the same equilibrium. Thus we can, without loss of generality, consider only the surface-driven symmetry breaking situation. In the rest of this article, the subscript *S* will refer to surface quantities and the subscript *B* refers to bulk quantities. In many cases, one of the two layers plays no role so we will omit this index when no confusion is possible. In the present example, in the surface-driven symmetry breaking situation, the bulk system plays no role at first order and we will thus denote for simplicity (*u*_*S*_, *v*_*S*_) *≡* (*u, v*). Similarly, the surface reaction function is denoted by ***f***_***S***_ *≡* (*f, g*) and *u* and *v* subscripts will denote partial differentiation.

##### Proposition 3.

*Under Assumption 1, let us consider a 1D-1D two-species model with surface concentrations* (*u, v*) *and such that*

- *the reaction functions* ***f***_***S***_ = (*f, g*) *satisfy the Turing conditions [45, Eqs. (2.31)];*
- *the bifurcation parameter δ ≡ d*_*v*_ *is the diffusion parameter of the v-species, which is the inhibitor, and the diffusion coefficient of the u-species (the activator) is constant and equal to d*_*u*_ = 1;
- *the coupling matrix* ***A*** *is diagonal with non-negative diagonal coefficients denoted by α, β, i*.*e*.

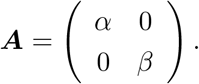

*Then ∂*_*δ*_|***P***_0_| *<* 0 *and the derivative of the bifurcation curve η ↦ δ*_*c*_(*η*) *at η* = 0 *is equal to*

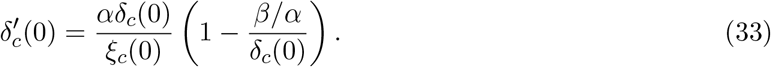

*Proof*. In this case, we have that

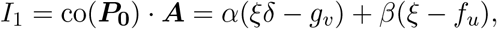

and

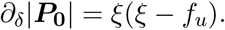

We recall [45, Eq. (2.25)] that the critical diffusion and critical wave number are linked by the relation

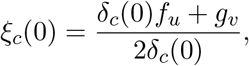

so that

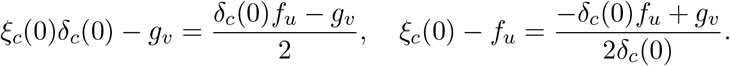

In particular, when the *u* and *v*-species are respectively the activator and the inhibitor, it follows that *f*_*u*_ *>* 0 and *g*_*v*_ *<* 0, and therefore *∂*_*δ*_|***P***_**0**_| *<* 0 and Eq. (28) simplifies to Eq. (33).

As a consequence, 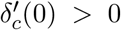 if and only if 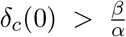. Since the classical Turing instability requires that the inhibitor diffusion coefficient is larger than the activator diffusion coefficient, and thus *δ*_*c*_(0) *≥* 1, this condition for *δ*_*c*_(0) is fulfilled when *β < α*. In other words, the prospect of patterning is reduced if the exchange rate of the activator is higher than the exchange rate of the inhibitor. This is always the case when *α* = *β* provided *δ*_*c*_(0) *>* 1. In order to enhance patterning it is sufficient to take *β > αδ*_*c*_(0), which can be understood as off-setting a small inhibitor diffusion with a high exchange rate. Note that the quantity *ξ*_*c*_(0) *− f*_*u*_ is the negative cofactor in the general case outlined in Section 4.2.1. This result is illustrated in Fig. 2 and Video 1 for the following Schnakenberg system [59]:

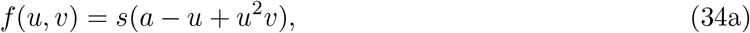

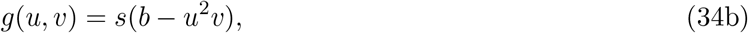

where 0 *< a < b* and *s >* 0.

#### 4.2.3 Chemotaxis systems and bulk-driven symmetry breaking

Let us consider now the bulk-driven symmetry breaking scenario with a population of chemotactic cells with concentration denoted by *c* which satisfies Eq. (5) in the bulk and two chemical species with equal equilibrium concentrations in the two layers. Similarly as before, we omit the *B* indexing since the surface does not play any role.

##### Proposition 4.

*Under Assumption 1 and the assumption that all the species have constant diffusion coefficients, let us consider the two following cases*.

1. *If only one species, the chemo-attractant with concentration u, diffuses through the interface and if the reaction functions satisfy f*_*u*_ *<* 0, *f*_*c*_ *>* 0 *and r*_*c*_ *<* 0 *(evaluated at the equilibirum point) then the chemo-attractant diffusion coefficient δ ≡ d*_*u*_ *is a bifurcation parameter for the uncoupled bulk system, i*.*e. there exists δ*_*c*_(0) *>* 0 *such that patterning occurs for δ < δ*_*c*_(0). *In the coupled case, if the exchange matrix* 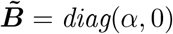 *is diagonal then*

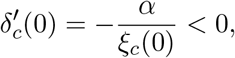

*and hence* 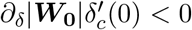 *so that coupling reduces the ability of the system to form patterns*.
2. *If the two reacting species* (*u, v*) *diffuse through the interface and satisfy Eqs*. (4)*-*(5) *with a reaction function* 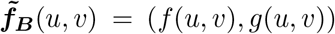 *and if the exchange matrix* ***B*** = *diag*(*α, β*) *is diagonal then the derivative at η* = 0 *of the bifurcation curve for δ ≡ d*_*v*_ *can be simplified using the following expressions:*

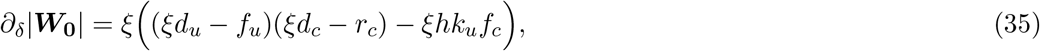

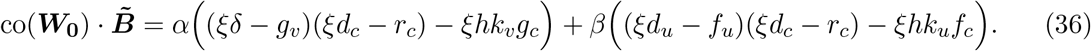

A typical example for the first case is the linear Keller-Segel model

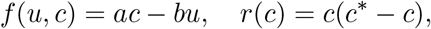

with coefficients *a, b, c*^*\**^ *>* 0.

1. *Proof*. 1. The linearized matrix of the bulk system reads,

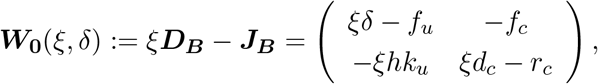

and its determinant is given by

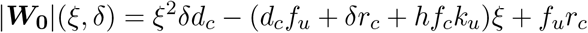

Since *f*_*u*_, *r*_*c*_ *<* 0, and thus *f*_*u*_*r*_*c*_ *>* 0, a necessary condition for this polynomial (in *ξ*) to take negative values for *ξ ≥* 0 is that its derivative in 0 is nonpositive, that is *δ <* (*hf*_*c*_*k*_*u*_ *−d*_*c*_|*f*_*u*_|)*/*|*r*_*c*_|. Then, the minimal value of this polynomial is given by *−*(*d*_*c*_*f*_*u*_ + *δr*_*c*_ + *hf*_*c*_*k*_*u*_)^2^*/*(4*δd*_*c*_) + *f*_*u*_*r*_*c*_. This function of *δ* is monotonically increasing between 0 and (*hf*_*c*_*k*_*u*_ *−d*_*c*_|*f*_*u*_|)*/*|*r*_*c*_| and is negative for *δ* smaller than a certain value which defines the critical value *δ*_*c*_(0) below which patterning occurs. Moreover, a direct computation shows that

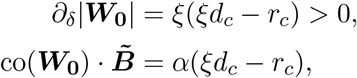

with diagonal matrix 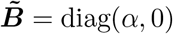. Consequently, Eq. (28) simplifies to

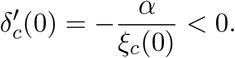
2. The full dispersion relation for the bulk reads

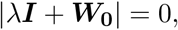

Where

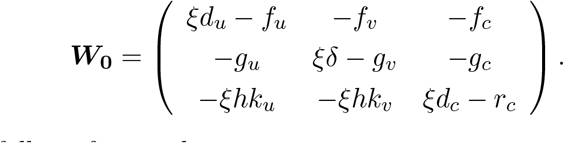

The conclusion thus follows from a direct computation. □ Although one can provide a definitive (negative) answer to the question of whether patterning is enhanced by the coupling in the first case, it seems impossible to draw general conclusions in the second case. Indeed it would require assessing the sign of the last expression Eq. (36), which, in a general setting, is difficult due to the number of degrees of freedom. However, one scenario where the equilibria of the reacting and diffusing species have the same values in the bulk and surface is where the cells are slave, with no feedback on the signalling molecules, so that *f*_*c*_ = *g*_*c*_ = 0. Further, the requirement of stability to homogeneous perturbations, i.e. when *ξ* = 0, gives *r*_*c*_ *<* 0, *f*_*u*_ +*g*_*v*_ *<* 0, and *f*_*u*_*g*_*v*_ *−g*_*u*_*f*_*v*_ *>* 0, with the latter two constituting standard Turing conditions. With the chemoattractant species, *v*, as the activator, and the assumption that *u, v* are a Turing pair in the absence of chemotaxis, so that *u* is the inhibitor, entails we additionally have *g*_*v*_ *>* 0 *> f*_*u*_ and hence *g*_*u*_*f*_*v*_ *< f*_*u*_*g*_*v*_ *<* 0. Thus the requirement that |***W***_**0**_| = 0 at the critical point gives

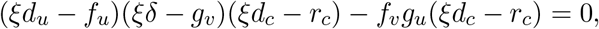

and hence the coefficient of *α* in co(***W***_**0**_) *·* 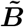 is given by

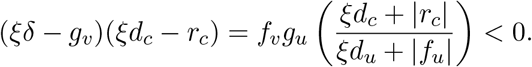

In contrast the coefficient of *β* is given by (*ξd*_*u*_ + |*f*_*u*_|)(*ξd*_*c*_ + |*r*_*c*_|) *>* 0. Hence, whether the bilayer enhances patterning depends on the between-layer transport of the inhibitor, *u* here, relative to that of the activator, *v* here, with sufficiently low relative activator between-layer transport acting to increase the ability to form patterns. We also note that the chemotactic parameters occur only in the grouping (*ξd*_*c*_ + |*r*_*c*_|), which factors out of both the coefficients of *α* and *β* so that, as might be expected given that the cells are slave to the signalling molecules, the chemotatic properties of the cells do not influence these observations.

### 4.3 Asymmetric equilibria in the 1D-1D model

In this section, we drop the assumption of the existence of global equilibria (Assumption 1) and we instead assume the following in the 1D-1D case.

**Assumption 2** (Asymmetric equilibria). *The equilibria of the uncoupled bulk and surface layers are different, i*.*e*.

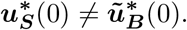

As usual, since the surface and bulk play a symmetric role in the 1D-1D case, we consider the surface-driven patterning case and in order to assess whether asymmetric equilibria have an influence on patterning, we focus on the contribution of the second term in Eq. (31).

#### Proposition 5.

*Under Assumption 2, it holds that*

*1. if* ***f***_***S***_ *and* ***f***_***B***_ *have continuous derivatives with invertible Jacobian matrices at η* = 0, *then there exists a differentiable equilibrium curve* 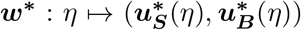 *defined on an interval* [0, *η*_0_) *with η*_0_ *>* 0 *and which satisfies Eqs*. (12)*-*(13);

*2. in this case, the derivative of the equilibrium curve at η* = 0 *is given by*

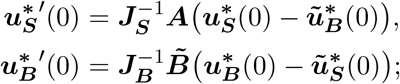

*3. consequently, the second term in Eq*. (31) *is equal to*

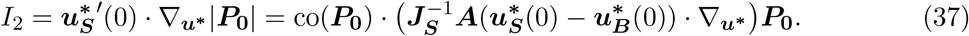

*Proof*. 1. This is a direct consequence of the implicit function theorem applied to the function

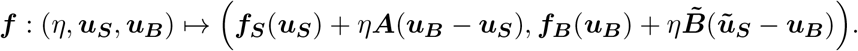

Indeed, this function vanishes at *η* = 0 by Assumption 2 and the determinant of the Jacobian matrix ***J***_***f***_ of ***f*** with respect to (***u***_***S***_, ***u***_***B***_) at *η* = 0 is given by

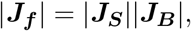

where ***J***_***S***_ and ***J***_***B***_ respectively denote the Jacobian matrices of the reaction functions of the surface and the bulk.

2. The derivatives of the equilibrium curves are computed by differentiating the relation

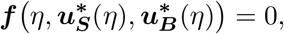

with respect to *η*.

3. The second expression of *I*_2_ comes from Jacobi’s formula for the differential of the determinant which implies that for a given vector ***p*** = (*p*_1_, …, *p*_*n*_)^T^,

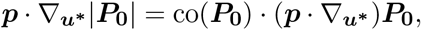

where we use the operator 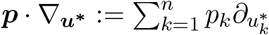 □

Since we only consider diffusion-driven instabilities, both Jacobian matrices ***J***_***S***_ and ***J***_***B***_ are invertible (as they must have eigenvalues with strictly negative real parts) and hence we are always in the situation where there exists a differentiable equilibrium curve.

From the second and third points of the previous proposition, we can conclude that the contribution of the asymmetry between equilibria on the first-order approximation of the bifurcation curve given by Eq.(28) is due to 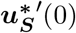 and it depends only on the difference between the equilibrium concentrations and on the reaction term of the surface but it does not involve the reaction term of the bulk layer. Thus, this contribution will dominate when the uncoupled equilibrium concentrations in the two layers are sufficiently different. This case is illustrated in Fig. 3. The situation is simpler in the strong coupling case, as explained in the forthcoming Section 5. but nonetheless progress can be made in simple examples, as we now show.

For example, we consider the Schnakenberg kinetics of Eq. (34), with *s* = 1, a coupling matrix ***A*** = ***B*** = diag(*α*, 0) and, to force a difference in the surface and bulk concentrations, we take the equivalent parameter for *a* in the bulk to be *a*_*B*_*≠ a*, with all parameters positive. The equilibria are given by the solution of the four simultaneous algebraic equations governing 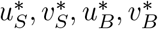, namely

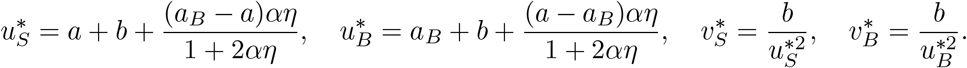

Thus

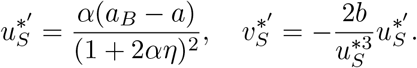

Hence, explicitly evaluating Eq. (37) with *d*_*u*_, *d*_*v*_ the diffusion coefficients of the respective species, we have

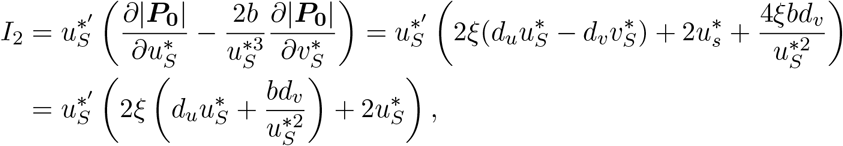

with the final equality arising from noting that 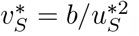. Hence, for this case we have

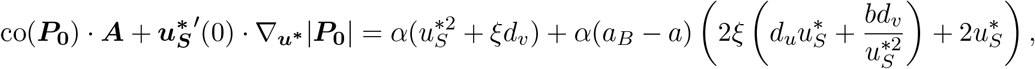

where we recall that this expression is evaluated at the critical point *η* = 0, *ξ* = *ξ*_*c*_(0), *d*_*v*_ = *δ*_*c*_(0) and 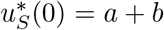. Thus, when *a*_*B*_ *> a*, this expression is always positive and the layering acts to inhibit pattern, whilst for *a > a*_*B*_ sufficiently large, we have that the difference between the resulting surface and bulk equilibrium concentration levels will enhance prospective patterning.

### 4.4 The case of the 1D-2D model

In this section, we consider exclusively the 1D-2D setting with a particular focus on bulk-driven symmetry breaking. In particular, the equilibria of the two uncoupled layers must be the same, that is Assumption 1 holds true.

#### 4.4.1 Large and small depth

In the 1D-2D case, the boundary condition Eq. (3) imposes a global homogeneous equilibrium identical in both layers (Assumption 1). Thus, for bulk-driven patterns, using (25), the formula Eq. (27) reduces to

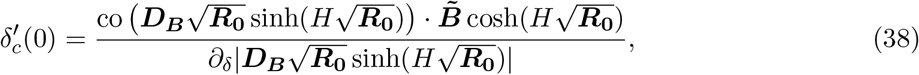

where 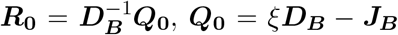 and the notation 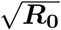 denotes any square root of ***R***_**0**_. We recall that

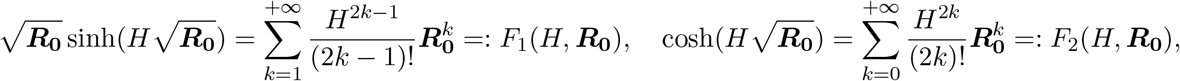

and thus the expression in Eq. (38) is a function of ***R***_**0**_ only. When *H →* 0, one can compute the following equivalent

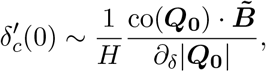

that is, up to a positive factor *H*^*−*1^, this is the same as in the 1D-1D case. This scaling is expected and consistent with the analysis for the reduction from 2D to 1D obtained when *H →* 0.

When *H* is large, we also expect that the bulk will pattern on its own without influence of the surface, that is 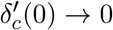. Using Jacobi’s formula and the series expansion in Eq. (22) we note that

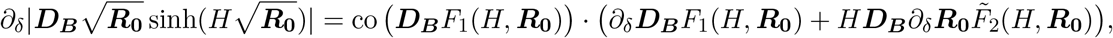

where

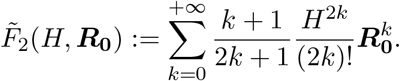

We can expect that, generically when *H* is large, the denominator in Eq. (38) will behave as *H* multi-plied by a quantity of the same order of magnitude as the numerator co 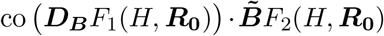. With this formal argument we thus expect that 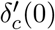 generically decreases as *H*^*−*1^.

#### 4.4.2 Two-species case

For a two-species system like the one in Section 4.2.2, we can make the above computation exact for any *H*. This follows from the fact that

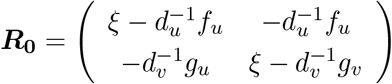

is nilpotent, satisfying 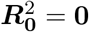 (for example by noting Tr(***R***_**0**_) = det(***R***_**0**_) = 0 at the critical bifurcation point *ξ* = *ξ*_*c*_(0), *d*_*v*_ = *δ*_*c*_(0), and then using the Cayley-Hamilton theorem). Further note that, in this case, the matrix ***R***_**0**_ has no square root. Thus it follows that,

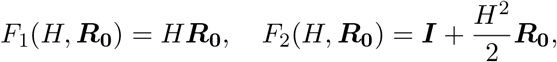

and Eq. (38) simplifies to

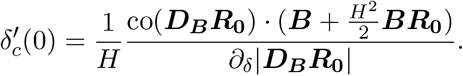

We note that since |***R***_**0**_| = 0,

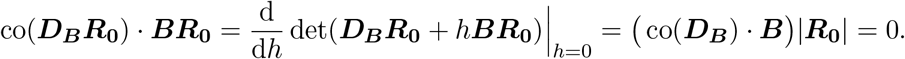

Moreover, ***D***_***B***_***R***_**0**_ = ***Q***_**0**_ so it follows that

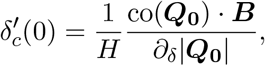

and this exact formula remains valid for all *H*.

##### Remark 2.

*It should be noted that the eigenvalues of* ***R***_**0**_ *at ξ* = *ξ*_*c*_(0) *and δ* = *δ*_*c*_(0) *are the solutions of*

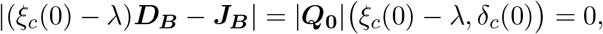

*so λ* = 0 *is always an eigenvalue of* ***R***_**0**_. *In the two-species case, this is the only eigenvalue since the polynomial ξ* ↦ |***Q***_**0**_| *(ξ, δ*_*c*_(0))*is of degree 2, vanishes at ξ*_*c*_(0) *and is non-negative in a neighbourhood of this root. Consequently*, ***R***_**0**_ *is a nilpotent matrix, as computed before. The same situation could possibly happen for an arbitrary, but even, number of species, although it would require some fine tuning of the parameters, in which case the functions F*_1_ *and F*_2_ *would actually be polynomials in* ***R***_**0**_. *However, for an odd number of species, the polynomial ξ ↦* |***Q***_**0**_| *(ξ, δ*_*c*_(0)) *is an even-degree polynomial and thus it must have at least one other root ξ*_0_ *< ξ*_*c*_(0). *Consequently, the matrix* ***R***_**0**_ *always has a positive eigenvalue λ* = *ξ*_*c*_(0) *− ξ*_0_ *>* 0 *so the functions F*_1_ *and F*_2_ *cannot be polynomials. This observation suggests that, in layered systems, 2-species models may not be representative of higher species models since the impact of nilpotency in 2-species models is a specific feature that may reflect a genuinely different behaviour compared to higher species models*.

#### 4.4.3 Patterning dynamics

Since Eq. (28) involves the characteristics of the patterning layer only, the critical bifurcation curve in the surface-patterning case for the 1D-2D system has the same first-order approximation as in the 1D-1D system. However, although the 1D-2D and 1D-1D systems look similar from the point of view of patterning conditions, the structure of the patterns is different depending on whether symmetry-breaking is driven by the surface or the bulk. As may be expected, in the former case, patterning preferentially occurs at the interface and is not observed to extend into the depth of a nonpatterning bulk system, at least for the cases we will consider below. For bulk-driven self-organisation, patterns can be observed throughout the bulk layer, with the potential of creating surface concentration inhomogeneities when these bulk patterns are formed close to the interface.

These phenomena are illustrated in Fig. 4 and Videos 2-3 for a reaction-diffusion-chemotaxis model (4)-(5) using the following piece-wise linear reaction functions: in the surface the reaction term is given by ***f***_***S***_(*u, v*) = (*f* (*u, v*), *g*(*u, v*)) with

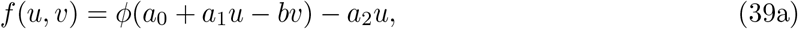

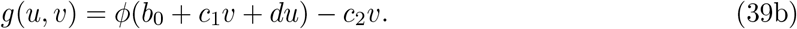

Here signalling molecules are produced by a fixed density of surface cells in an autocatlytic feedback response, with the continuous piece-wise linear function *ϕ*(*ζ*) = max(0, min(*ζ, M*)) for a given *M >* 0, and parameters *a*_0_, *a*_1_, *b, a*_2_, *b*_0_, *c*_1_, *d, c*_2_ *≥* 0. In this definition the value *M* is chosen sufficiently large so that the reaction function behaves as the identity function around the equilibrium value (obtained by solving the linear system, with *ϕ* taken as the identity). The threshold values 0 and *M* are added to ensure the positivity of the solutions and to prevent blow-up. This model has been introduced in the seminal work [30]. In the bulk we consider the same system but where the chemical species are produced by the chemotactic cells without feedback, so that ***f***_***B***_(*u, v, c*) = (*f* (*u, v, c*), *g*(*u, v, c*), *r*(*c*)) with

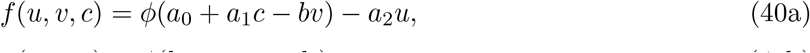

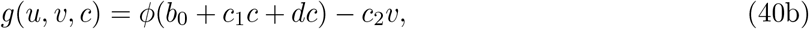

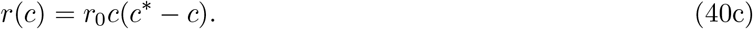

The parameters in both systems and the equilibrium cell concentration *c*^*\**^ are chosen so that 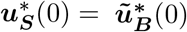 For a given set of parameters, the critical diffusion of the *v*-species in the surface and the bulk are, respectively, denoted in the caption of the figures by 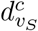and 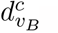.

**Figure 4:**
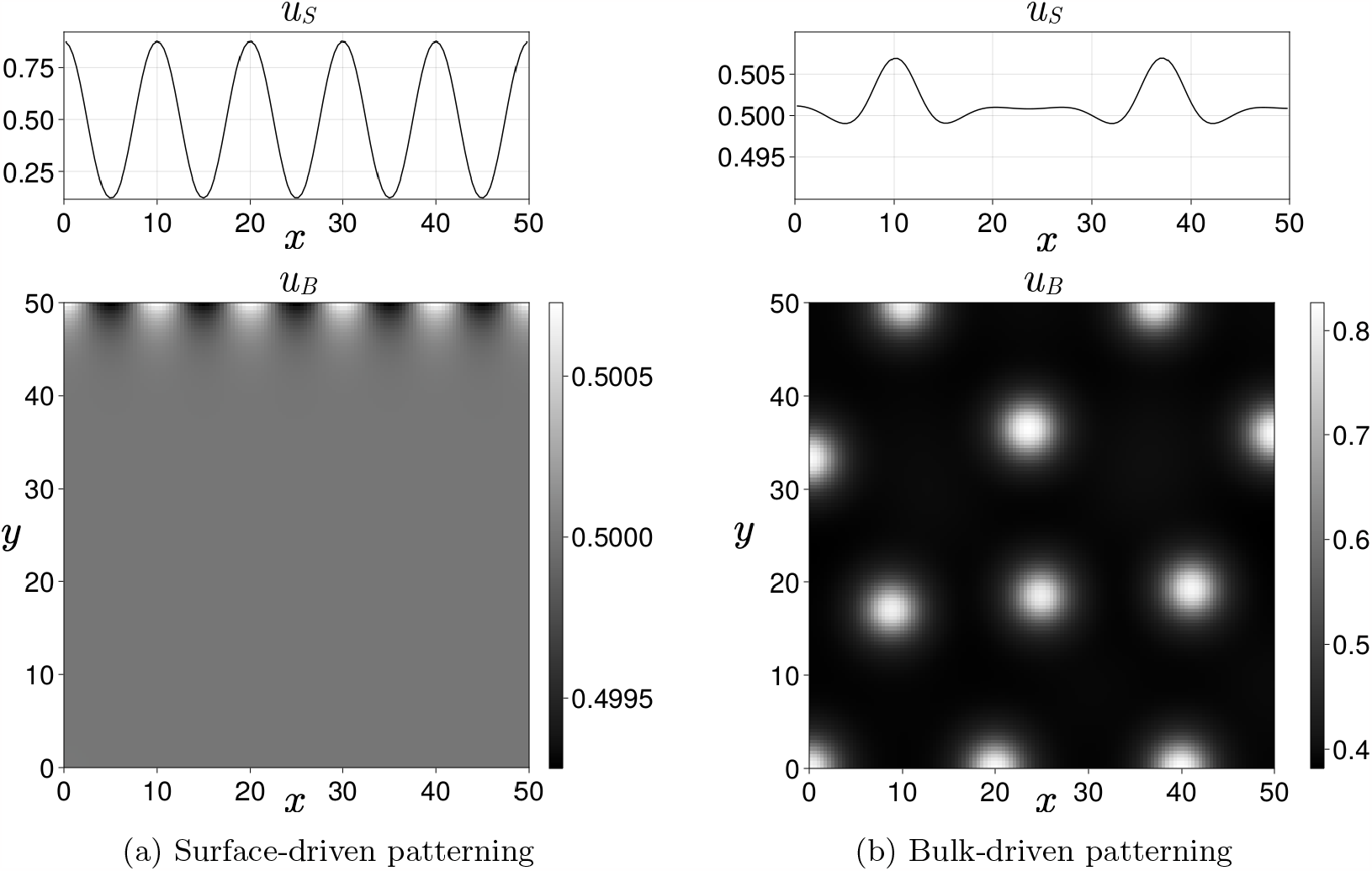
Surface and bulk patterning with *η* = 0.01 in the pseudo-linear 1D-2D system (39)-(40) computed in a 128 *×* 128 grid at time *t* = 500 after a steady state is reached. Surface parameters: *a*_0_ = 1, *a*_1_ = 2, *b* = 2, *a*_2_ = 2, *b*_0_ = 0.25, *c*_1_ = 1, *d* = 1, *c*_2_ = 2. Bulk parameters: *r*_0_ = 0.2, *a*_0_ = 3, *a*_1_ = 8, *b* = 2, *a*_2_ = 7, *b*_0_ = 0.75, *c*_1_ = 0.5, *c*_2_ = 1.5, *d* = 1, *c*^*\**^ = 0.25, *χ* = 50. (a) Surface-driven patterning: 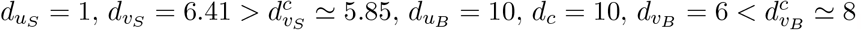 . (b) Bulk-driven patterning: 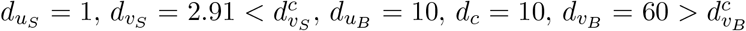 . Note that for both plots the original *y*-coordinate is used, with the no-flux condition at *y* = 0 and the surface at *y* = *H* = 50. A brief description of the numerical method used can be found in Appendix A as well as a link to the numerical code freely available online.

In this situation, since one layer is in a patterning state, spot patterns appear simultaneously throughout the patterning layer and propagate through the interface to the non-patterning layer. The amplitude of the patterns created in the non-patterning layer depends on the coupling strength *η*, and it has been confirmed numerically that they are directly proportional (results not shown). In particular, since we consider the case of small *η* in all of the numerical simulations presented in this section, the amplitude of the patterns in the non-patterning layer is typically much lower than in the patterning layer. For instance in Fig. 4b and Fig. 5, the amplitude of the surface patterns is about one percent or less of the value of the homogeneous steady state, but it would be proportionally larger for larger values of the coupling strength *η* (in the cases shown, we took *η* = 0.01). The same is true for the bulk amplitude in Fig. 4a.

**Figure 5:**
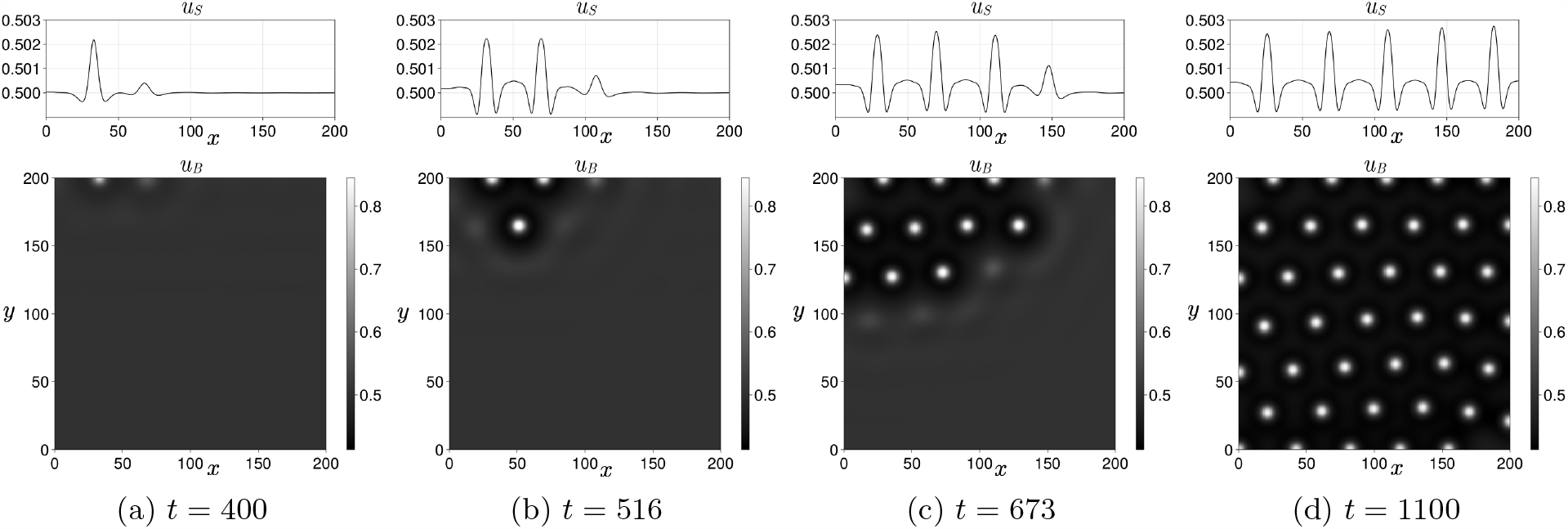
Wave pattern with *η* = 0.01 computed on a 512 *×* 512 grid until a steady state is reached. Surface parameters: 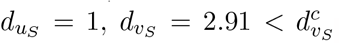 . Bulk parameters: *χ* = 120, 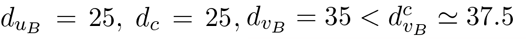, *β*_*B*_ = 180. The other parameters are the same as in Fig. 4.

The patterning dynamics may be different when the patterns are created by coupling two nonpatterning layers with sufficiently large and different exchange rates, i.e. by taking ***A*** or ***B*** sufficiently different from the identity, similarly to the system in Section 4.2.2. and Fig. 2b. In particular, Fig. 5 shows the result for the piece-wise linear system (39)-(40) with ***A*** = ***I*** and ***B*** = diag(1, *β*_*B*_). Starting from a random perturbation of the homogeneous equilibrium, when the exchange rate *β*_*B*_ of the inhibitor *v* is sufficiently large, patterns appear locally at the interface and then propagate throughout the bulk and the surface (see also Video 4).

The pattern propagation in the bulk (Fig. 5) can be understood as a multi-stability phenomenon: although linearly stable, the homogeneous state in the bulk is only one stable equilibrium of the PDE, and patterns in the uncoupled system can be obtained by choosing an initial condition very far from the homogeneous equilibrium. In the simulation, the initial condition is a small random perturbation of the homogeneous state and the bulk independently thus does not typically escape from this linearly stable state. However, when coupling is added, the interface plays the role of an active boundary able to locally drive the system far from the homogeneous equilibrium before spreading throughout the domain, in a similar fashion as the models introduced by [39, 47].

## 5 Strong coupling in the 1D-1D case

### 5.1 Asymptotic reduction

When *η ≥* 1 and for sufficiently nice ***A*** and ***B***, the concentrations ũ_***B***_ and ***u***_***S***_ are expected to converge towards a common value ***u***. For simplicity, we consider here only the case *m* = *n*, but the extension to the general case would be straightforward. Thus, splitting ***u***_***S***_ and ***u***_***B***_ as

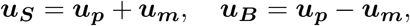

Where

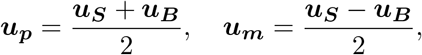

provided that ***u***_***S***_ and ***u***_***B***_ remain uniformly bounded, the half difference ***u***_***m***_ satisfies

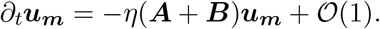

As a consequence, if ***A*** and ***B*** are such that their sum ***A*** + ***B*** has only non-negative eigenvalues (for instance when ***A*** and ***B*** are both symmetric and positive definite), it follows that ***u***_***m***_ = *O*(*η*^*−*1^). Up to an error of order *η*^*−*1^, the concentrations ***u***_***S***_ and ***u***_***B***_ are thus equal to their average ***u***_***p***_. The dynamics of ***u***_***p***_ can be found by multiplying Eq. (10) by ***A***^*−*1^ and Eq. (11) by ***B***^*−*1^ and summing the two resulting equations so that the exchange term cancels out. Again, up to an error of order *η*^*−*1^, the behaviour of ***u***_***p***_ is given by the following equation

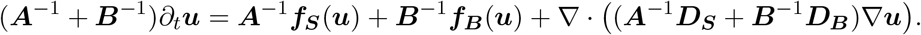

For instance, in the simple case where ***A*** = ***B***, corresponding to no sinks or sources at the interface, ***u*** satisfies

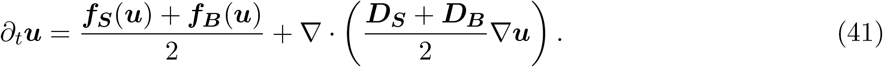

A further case with a simple reduction occurs when ***f***_***S***_ = ***f***_***B***_ so that the reaction term remains unchanged while the diffusion matrix becomes a linear combination of the surface and bulk diffusion matrices. In addition, for a mass conserving system with ***f***_***S***_ = *−****f***_***B***_, the reduced equation Eq. (41) simply becomes a pure (cross-)diffusion. However, in full generality, the behaviour of the solution does not readily simplify, since the properties of the new reaction term cannot be expected to immediately follow from the properties of the surface and bulk reaction terms.

### 5.2 Patterning conditions for two-species systems

In the case of two coupled two-species reaction-diffusion systems with 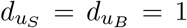, the critical bifurcation parameter of the inhibitor in the surface can be simply computed from Eq. (41) and is given by

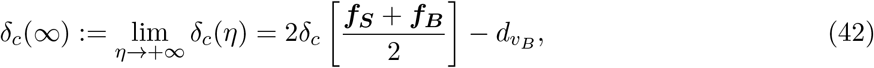

where we denote by *δ*_*c*_[***f***] the critical inhibitor diffusion of a reaction-diffusion system with reaction term ***f*** = (*f, g*) and activator diffusion *d*_*u*_ = 1. Note that there is a balance between the diffusion coefficients of the two layers. In particular, and as in the small coupling case (Section 4.2.1), the critical diffusion of the *v*-species in the surface in the coupled system can be made as small as desired, provided that the diffusion in bulk 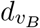 is large enough.

For later convenience, we recall the following formula corresponding to a standard reaction-diffusion system with ***f*** = (*f, g*) [45, Eq. (2.27)]:

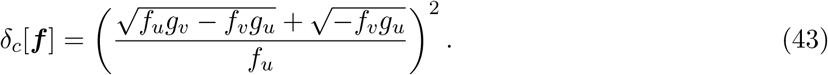

#### 5.2.1 Stabilization by coupling

When the two coupled systems are identical, ***f***_***S***_ = ***f***_***B***_, with a common critical bifurcation parameter for the inihibitor diffusion denoted by *δ*_*c*_(0), then, asymptotically when *η* is large,

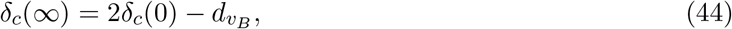

and in particular, if the bulk is in a non-patterning state, i.e. 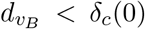, then *δ*_*c*_(*∞*) *> δ*_*c*_(0). As might be expected, and similarly to the small coupling case (Section 4.2.), this means that a large coupling has a stabilizing effect: a larger diffusion of the inhibitor in the surface is needed to de-stabilize the homogeneous state. This observation is illustrated in Fig. 6a.

**Figure 6:**
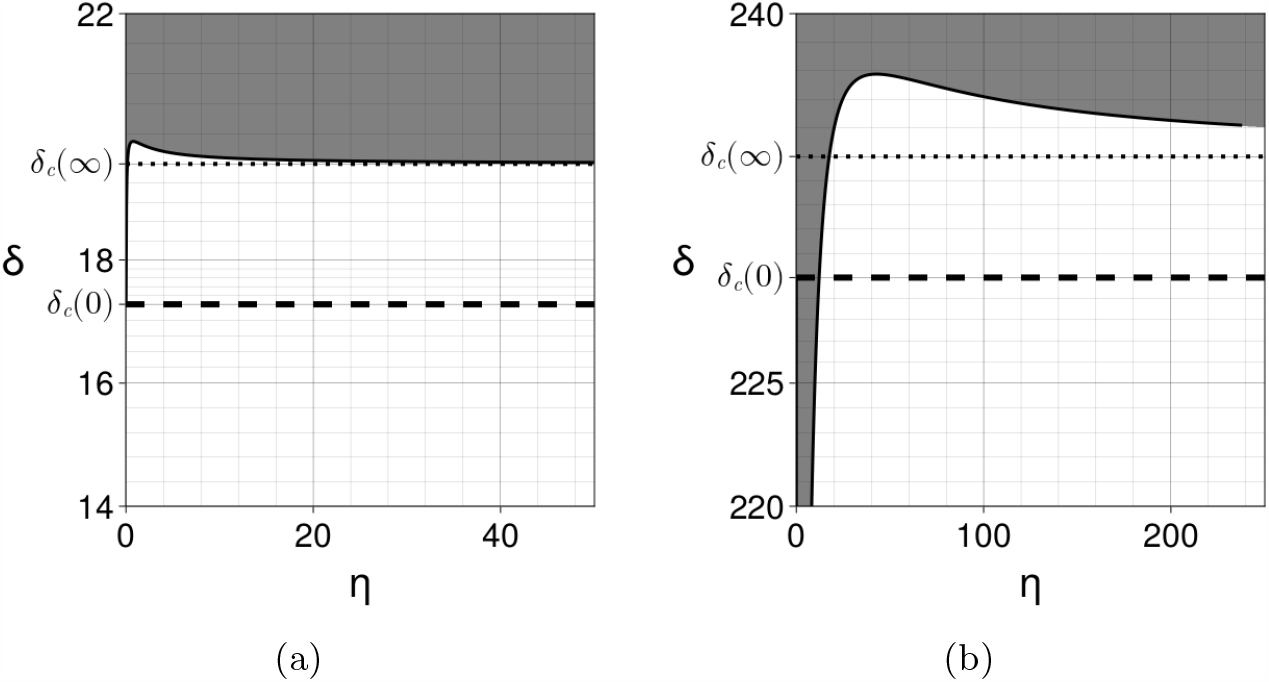
Large coupling asymptotics. The dotted horizontal line indicates the asymptotic value *δ*_*c*_(*∞*) computed using Eq. (42). (a) Coupling two identical Schnakenberg systems with the same parameters as in Fig. 2. The asymptotic value of the critical bifurcation parameter is larger than *δ*_*c*_(0). (b) Coupling two pseudo-linear systems with Jacobians given by Eq. (46) with *p* = 2 for the bulk and *p* = 200 for the surface. The bulk is in a patterning state with diffusion coefficients 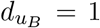 and 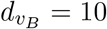. Consequently, for small *η* the coupled system is in a patterning state for all values of 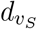 . However, since *δ*_*c*_(*∞*) *> δ*_*c*_(0), for *η* sufficiently large and 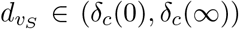, the coupled system is in a non-patterning state and thus the coupling stabilizes the homogeneous equilibrium. See also Video 5.

More generally, the analysis presented in Section 4 reveals that if a layer is in a patterning state, then for small coupling, the coupled system remains in a patterning state. This can be seen as a consequence of the fact that the derivative 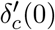 of the critical parameter is an intrinsic property of the considered layer. Consequently, if *δ* is chosen above the critical value *δ*_*c*_(0) and 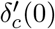, then the coupled system must remain in a patterning state for *η* sufficiently small (when 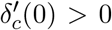 and *δ* is close to *δ*_*c*_(0), a first order approximation of this value would be 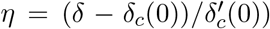. Therefore, if one wants to use the coupling to stabilize a homogeneous state, it is necessary to consider a sufficiently large coupling strength. When the two layers are independently in a patterning state, i. e. when 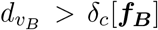 and 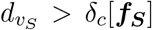, an asymptotically non-patterning state for the coupled system corresponds to the case where

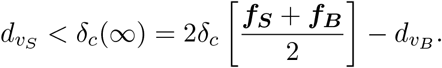

It is possible to find such parameters 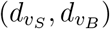 if and only if

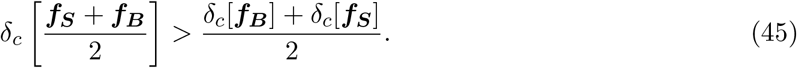

This strict midpoint-concavity property requires that ***f***_***B***_ ***f***_***S***_. Checking this property for two arbitrary reaction functions heavily depends on the form of these functions. When the reaction function depends linearly on its parameters, using the explicit formula Eq. (43) for the critical diffusion, checking Eq. (45) reduces to checking the convexity properties of this several-variable function (note that midpoint convexity is equivalent to convexity for continuous functions). For instance, it is straightforward to select reaction functions which satisfy this concavity property: one can consider a family of (pseudo)-linear models of the form (39) parametrised by a single parameter *p >* 1 such that the Jacobian matrix of the equilibrium system is given by

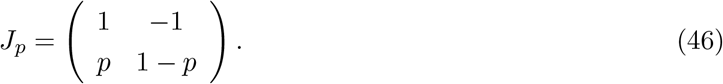

In this case

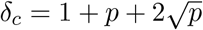

is a concave function of *p*. This situation is illustrated in Fig. 6b and Video 5.

#### 5.2.2 Strong coupling patterns

Reciprocally, if 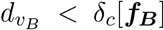 and 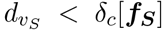 (i.e. the two layers are independently in a nonpatterning state), then the homogeneous state of the coupled system, with *η* large, is unstable whenever 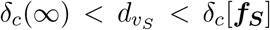. Using Eq. (42), such a value of 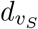 will exist if and only if the following midpoint-convexity condition is satisfied:

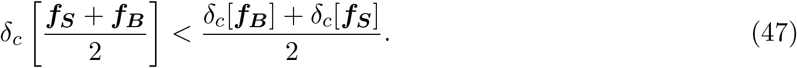

Once again, it is necessary to consider two different systems in the surface and bulk layers. Note that in contrast to the small coupling case (Section 4.3.), the patterning condition (47) is typically more straightforward to ascertain, as it reduces to computing the convexity properties of a function. For instance, the condition (47) is always satisfied for the Schnakenberg (34) [59] and Gierer-Meinhardt [23], [45, Eq. (2.8)] systems, as it can be checked using the formula (43). An example is shown in Fig. 3b and Video 6.

## 6 Beyond the asymptotic cases

For intermediate values of *η* many different scenarios are possible. An important point to note is that the Turing conditions for each layer at *η* = 0 do not ensure that the coupled system at *η >* 0 is stable without diffusion (i.e. that *ξ* = 0 is a stable mode). For instance, taking two identical 1D-1D layers with Jacobian matrix ***J*** and ***A*** = ***B***, as in Eq. (16), the stability of the mode *ξ* = 0 can be checked by computing the eigenvalues of the block matrix

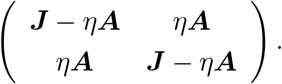

Using the formula for block-diagonal determinants, the eigenvalues of this matrix are the eigenvalues of the matrix ***J*** and the eigenvalues of the matrix ***J*** *−* 2*η****A***. If ***A****≠* ***I***, then the latter may have non-negative eigenvalues, indicating that the mode *ξ* = 0 is unstable. For instance for 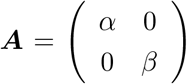, the mode *ξ* = 0 is unstable for all *η* between the roots of the polynomial (in *η*)

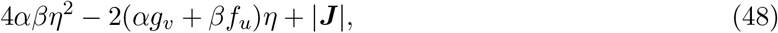

as soon as this polynomial has two real roots. This phenomenon is illustrated on Fig. 7.

**Figure 7:**
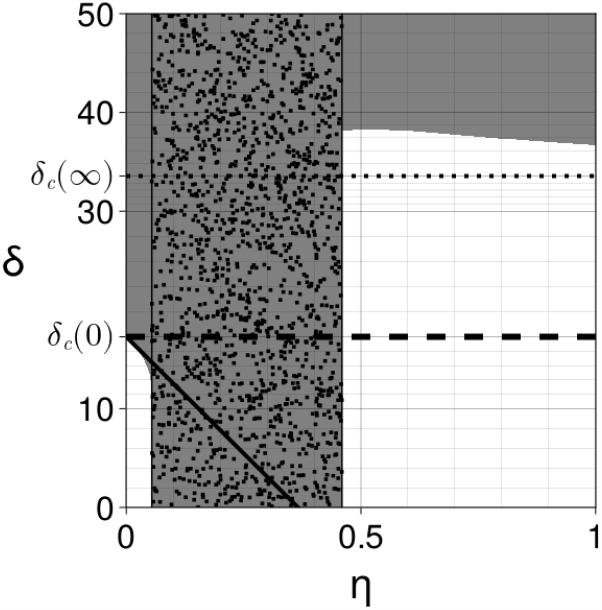
Coupling two identical Schnakenberg systems with *a* = 0.2305, *b* = 0.7695, *s* = 2, 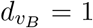 and ***A*** = ***B*** diagonal with coefficients *α* = 1 and *β* = 40. The behaviour of the boundary of the instability region (in grey) near *η* = 0 and for large *η* is well-predicted respectively by Eq. (33) (solid thick line) and Eq. (42) (dotted horizontal line). For *η* between the roots of the polynomial Eq. (48) (depicted by the randomly dotted band), the 0-th mode is unstable, which indicates a non-Turing type instability.

## 7 Conclusion and discussion

During development, many organs are characterised by a bilayer geometry, such as the skin and internal epithelium. However, the precise effect of this bilayer coupling on self-organisation dynamics is underexplored. In this study, we have investigated patterning conditions for bilayer reaction-cross-diffusion systems with weak and strong coupling.

Our analysis provides a quantitative description of this coupling effect and shows in particular that not only can spatial patterns emerge from the coupling of two non-patterning layers, but also that the coupling of two independent patterning layers can stabilize a homogeneous equilibrium and thus diminish patterning. Furthermore, the reaction-cross-diffusion systems investigated in this study show that the classical paradigm of local activation and long-range inhibition can be weakened by considering alternative mechanisms specific to the bilayer geometry. In particular, we found that different transport rates between the two layers, or the asymmetry of the equilibrium states of the model components between the two layers, can have a critical influence on pattern formation. In the asymptotic regimes of weak and strong coupling, we proved that these constraints are necessary to induce patterning between two non-patterning layers and we highlighted several explicit examples where they are sufficient. Furthermore, we also classified the cases where the coupling has a negative effect on patterning when one or both layers is independently in a patterning state. We observed an additional complication that the layered coupling can disrupt the stability of the homogeneous steady state that is required for Turing’s mechanism, further highlighting the need to consider the coupled system rather than layers in isolation. Further behaviours observed in these bilayer systems include a wave of patterning induction, contrasting the behaviour of the classical Turing system, which patterns via simultaneous self-organisation across the domain. In particular, the combined wave and patterning dynamics bears some resemblance to cell density waves seen in feather patterning. We note that feather placode formation is also typically accompanied by an additional signalling-molecule wave [28], so this phenomenological observation needs more careful study.

More generally, our analysis covers the cases of 1D-1D and 1D-2D systems and these findings offer the prospect of improving our understanding of patterning mechanisms under the control of signalling molecules, and responses to them, in organs incorporating a bilayer structure in their development, such as the skin. Other organs that exhibit an analogous bilayering alongside spatial patterning in development include the villi of the gut, and the branches of the lung and various models for these systems are reviewed in [1, 42].

In addition to investigating this application to biological systems in detail, several other extensions of this work could be considered. First of all, the models studied in this paper are one-dimensional along the *x*-axis, as Turing patterning conditions are independent of the dimension, modulo geometric effects on wavemode selection [32]. However, in order to quantitatively study the patterning dynamics or to develop the analysis of pattern types, the models should be extended to a 2D surface and a 2D or 3D bulk. This configuration would also be more realistic from the perspective of prospective applications, such as feather array patterning [28]. From a mathematical perspective, and following the seminal developments of Shaw et. al [60] and Ermentrout [17], one should not only study the emergence of patterning but also the self-organized pattern shapes, requiring weakly-nonlinear bifurcation analysis. Weakly nonlinear analysis in the 1D-1D setting has been done in some special cases, e.g. [9, 8], so extending the present theory to this case would be challenging but plausible. In the 1D-2D setting, however, substantial new mathematics would need to be developed to perform such analysis given the complexity of the eigenfunctions in this setting. Another mathematical direction meriting further investigation is a consideration of Hopf/Wave bifurcations in these coupled systems, whereby linear instability leads to spatiotemporally oscillating solutions. Such instabilities are not possible in the classical two-species case, but exist in three-species models or models incorporating inertial (hyperbolic) effects [32, 57]. Regarding potential biological applications, see, for example, [10].

Furthermore, when the bulk has a non-zero depth along the *y*-axis, another generalisation could consider when the two layers do not admit the same equilibrium state. This would lead to a nonhomogeneous equilibrium state, constant along the *x*-axis of the interface, but possessing a gradient along the *y*-axis. Studying the linear stability of such a system would require perturbing about a heterogeneous steady state and, for instance, may further localise patterning to near the interface, in contrast to the situation obtained in Fig. 5. We remark that even in simple 1D scenarios, spatial heterogeneity often requires the study of specific asymptotic regimes to make progress [22]. Another direction is to consider the influence of the interface shape and mechanical structure on patterning. For instance, a fundamental question is whether a local deformation of the interface could have an influence on patterning, noting that the self-organisation is associated with a local mechanical compression and deformation of the epithelium [25, 28], as well as motivating earlier and more recent theoretical work [11, 13].

In summary, we have derived conditions for self organisation for a coupled bilayer system, with reaction-cross diffusion dynamics in each layer, under a number of limiting cases, such as thin layers and weak or strong coupling. The exploration of these conditions has revealed how the bilayering influences pattern formation, and can act to enhance or inhibit the prospect of patterning. In particular, our study emphasises that a detailed comparison of theory with observation for developmental periodic-structure formation, or chemical patterning in layered reaction diffusion systems [14], has to accommodate the prospect that considering the layers in isolation is insufficient to determine the presence, or absence, of self-organisation.

## Acknowledgments

The authors are grateful to Denis Headon for his advice and for the useful discussions regarding the biology of the skin. AD and SSL acknowledge the hospitality of the Wolfson Centre for Mathematical Biology of the University of Oxford (UK) where part of this research was carried out. This work is partially funded by the following grants: KAKENHI Transformative Research A (Grant number: JP22H05110) and KAKENHI Fostering Joint International Research (Grant number: JP17KK0094).

## A Numerical methods

The code used to produce all the simulations and plots presented in this article is freely accessible at

https://github.com/antoinediez/BilayerReactionCrossDiffusion

It is written in the `Julia` programming language and is based on the following open source libraries.

- All the plots and videos are produced using the `Makie.jl` data visualization ecosystem developed by [12].
- The bifurcation curves and instability regions in the (*η, δ*) plane for the 1D-1D case are computed using the Routh-Hurwitz criterion and the explicit equations Eqs. (20)-(21). Once the reaction terms have been specified by a formula, all the computations which define the dispersion relations and the Routh-Hurwitz criterion only involve differentiation and polynomial calculus so they can be done entirely symbolically. For this, we use the `Symbolics.jl` package developed by [27]. In order to compute the bifurcation curves, we compute the numerical solution of the dispersion relation using the nonlinear solver `NLsolve.jl` developed by [43]. The advantage of this symbolic approach is that the code is entirely independent of the particular form of the reaction functions. Running the code then only requires the user to specify the reaction functions by their symbolic formula.
- In the code provided, a model (i.e. a set of reaction functions) can be defined as a Julia function using the syntax detailed in the script `TuringSpace/models.jl`. The Turing space can then be computed automatically by running the script `TuringSpace/main.jl` after specifying the set of modes to investigate and the desired bifurcation parameters.
- The simulations of the 1D-1D and 1D-2D systems are implemented using a classical method of lines with an upwind scheme for chemotaxis [29] and an implicit adaptive time-stepping using the ODE solver `DifferentialEquations.jl` developed by [54].

More precisely, the problem is classically discretized in space on a grid of size *N*_*x*_ *×N*_*y*_ in 2D and of size *N*_*x*_ in 1D. In our simulations, we chose *N*_*x*_ = *N*_*y*_ with a value ranging from 56 to 256 (for the different simulations and values, we have checked that the results shown do not depend on the discretization size). The Laplacian is discretized on each grid cell using a standard central differences five-point stencil discretization in 2D and a three-point discretization in 1D. For the chemotaxis part, using a cell-centered finite volume approach, the space derivative along each dimension is computed as a flux difference where the flux at each cell boundary is computed via a third order upwind scheme [29, Section I.3]. Once all the spatial derivatives and reaction terms in each cell have been computed, we obtain a high-dimensional ODE system (of size *N*_*x*_ *× N*_*x*_ *× N*_*y*_ in the 1D-2D case) in time which is solved using an implicit adaptive Euler scheme. In order to reduce the computation time, a sparsity pattern associated to this highdimensional system is computed automatically beforehand, again using the routines implemented in `DifferentialEquations.jl`.

Each simulation presented in this article can be reproduced using the code provided by running the script `BilayerRCD/main.jl`, potentially after changing the parameters which are indicated in the caption of the figure corresponding to this simulation.

## B Supplementary videos

This article is supplemented with the following illustrative videos which show example simulations of 1D-1D and 1D-2D reaction-diffusion-chemotaxis systems. These videos can be accessed online along with the numerical code to reproduce them at

https://github.com/antoinediez/BilayerReactionCrossDiffusion

**Video 1**. Example of patterning with small coupling for the system described in Fig. 2b with *η* = 0.01 and 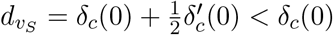.

**Video 2**. Example of surface-driven patterning for the 1D-2D system described in Fig. 4a.

**Video 3**. Example of bulk-driven patterning for the 1D-2D system described in Fig. 4b.

**Video 4**. Wave patterning in the 1D-2D system described in Fig. 5.

**Video 5**. Example of stabilization of a homogeneous equilibrium using a large coupling in the system described in Fig. 6b with *η* = 200 and 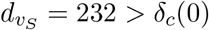.

**Video 6**. Example of patterning with large coupling for the system described in Fig. 3b with *η* = 100 and 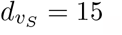.

## Footnotes

1 We suppress the dependence on time, *t*, for notational convenience.

